# SIANN: Strain Identification by Alignment to Near Neighbors

**DOI:** 10.1101/001727

**Authors:** Samuel S. Minot, Stephen D. Turner, Krista L. Ternus, Dana R. Kadavy

## Abstract

Next-generation sequencing is increasingly being used to study samples composed of mixtures of organisms, such as in clinical applications where the presence of a pathogen at very low abundance may be highly important. We present an analytical method (SIANN: Strain Identification by Alignment to Near Neighbors) specifically designed to rapidly detect a set of target organisms in mixed samples that achieves a high degree of species- and strain-specificity by aligning short sequence reads to the genomes of near neighbor organisms, as well as that of the target. Empirical benchmarking alongside the current state-of-the-art methods shows an extremely high Positive Predictive Value, even at very low abundances of the target organism in a mixed sample. SIANN is available as an Illumina BaseSpace app, as well as through Signature Science, LLC. SIANN results are presented in a streamlined report designed to be comprehensible to the non-specialist user, providing a powerful tool for rapid species detection in a mixed sample. By focusing on a set of (customizable) target organisms and their near neighbors, SIANN can operate quickly and with low computational requirements while delivering highly accurate results.

## Introduction

There are many different methods that characterize the mixture of organisms present within a metagenomic dataset. Such datasets are generated when a complex environmental sample is processed by a “next-generation” high-throughput genome sequencing protocol, and they consist of large numbers of short nucleotide sequences. Each sequence represents a small fragment of a randomly selected genome from the very large collection of genomes present in the source sample. Those sequences indicate the presence of one organism or another according to their similarity to a set of known reference genomes. While a given sequence may be unique to one species, it also may be found in diverse organisms across the tree of life. Therefore, one analytical challenge (among many) is to take that collection of sequences (likely numbering in the millions) and accurately determine what species are present in the sample. Here we describe a novel method (SIANN: Strain Identification by Alignment to Near Neighbors) that is specifically designed to rapidly detect a set of targeted organisms from a metagenomic dataset by aligning reads to genomic regions that are unique at the strain or species level.

The analytical question motivating a particular piece of metagenomic bioinformatic analysis may vary widely by user and sample type (Segata, et al., 2013). For example, the function of the human gut microbiome may depend on the relative abundance of hundreds of species of bacteria and the types of metabolic genes they contain (Wu, et al., 2011; Schloissnig, et al., 2013). In contrast, the clinical treatment of a patient may depend on whether or not a particular virus, or a consortium of co-infecting pathogens, is/are detected in their blood. It is this second class of presence/absence questions that SIANN is designed to address. SIANN is appropriate for situations in which a user wants to know whether a particular organism or set of organisms is present in a sample, but isn’t interested in the functions encoded in their genomes, the relative abundance of each organism, or any other more in-depth analysis.

## Methods

### Approach

Metagenomic classification methods are based on a wide variety of theoretical underpinnings. The basic varieties include alignment of reads to various nucleotide databases or exact matching to nucleotide or protein signature sequences (or *kmers*). A representative set of recent methods are described in Table 1 (also see Bazinet & Cummings 2012).

**Table 1.** Summary of methods for metagenomic classification.

| Name | Method | Reference |
| --- | --- | --- |
| **MEGAN** | Alignment to large nucleotide database | Huson, et al., 2011 |
| **PhymmBL** | Alignment to large nucleotide database with interpolated Markov models | Brady & Salzberg, 2011 |
| **Metaphyler** | Alignment to clade-specific marker genes | Liu, et al., 2011 |
| **MetaPhlAn** | Alignment to clade-specific marker genes | Segata, et al., 2012 |
| **LMAT** | Nucleotide kmer matching | Ames, et al., 2013 |
| **Kraken** | Nucleotide kmer matching | Wood & Salzberg, *in submission* |
| **Sequedex** | Protein kmer matching | Berendzen, et al., 2012 |
| **mOTU** | Alignment to universal marker genes | Sunagawa, et al., 2013 |
| **Phylosift** | Insertion into reference nucleotide and protein alignments | Darling, et al., *in preparation* |

Overall, these methods are designed to either classify individual reads to, and/or predict the total abundance of, clades (*e.g.* genus or species) across the entire tree of life. They generally require reference databases that are very large and/or require a large amount of processing to generate. The gap SIANN is designed to fill is when the entire tree of life is irrelevant, and only predefined subsets of organisms need to be detected. For an underlying method we chose read alignment to diagnostic genomic regions because the algorithms for read alignment are highly parallelizable and have been optimized heavily by the community at large (the current implementation of SIANN uses bowtie2 [Langmead & Salzberg, 2012] for the alignment function, but can be adapted to any alignment algorithm). This approach is distinct from using clade-specific marker genes (Segata, et al., 2012) because unique regions that are larger, smaller, or outside of genes can also be used. Furthermore, this approach supports the rapid construction of custom databases using reference genome sets that require only minimal user-supplied structure.

To understand the principle at work, consider a set of reads that have been aligned to the genomes of several strains belonging to two species. Some regions of those genomes are species-specific, some are strain-specific, and some are shared (Figure 1a). When a set of reads is aligned to those genomes such that each read is placed in as many locations as it has a match (at a reasonably stringent threshold), visual inspection of the distribution of reads yields an intuitive understanding of the true source organism as Species I/Strain B (Figure 1b). If Strain B were not present in the reference database, it would still be clear that the organism was an unknown strain of Species I.

**Figure 1.**
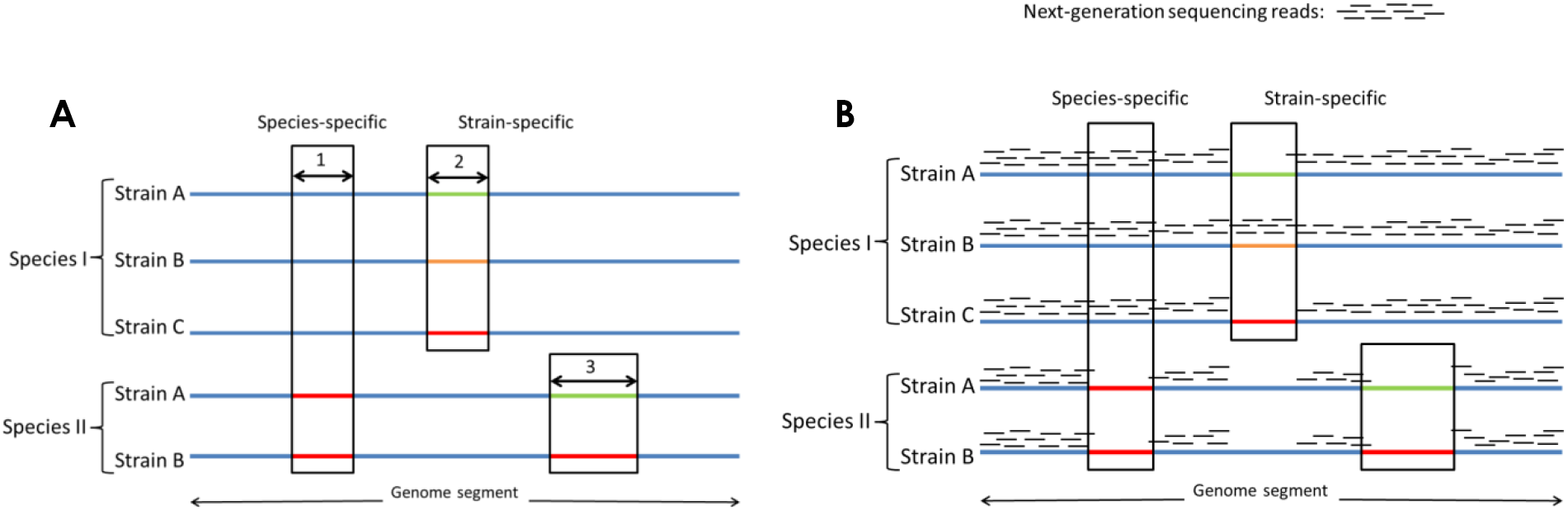
A) For a group of strains belonging to two different species, some regions may be unique to each species (region 1), while other regions may be unique to strains within each species (regions 2 and 3). B) A set of reads are aligned to these genomes, and the ones that align in a species- or strain-specific manner are identified by the combination of genomes to which they align. In this example, Strain B of Species I is the organism identified.

The unique identification of a species or strain is quantified by the proportion of the genome that is determined to be species- or strain-specific (defined as reads that are aligned to regions that are species- or strain-specific). Each species and strain is then assigned a numerical measure of the proportion that is covered by these diagnostic reads, and that proportional measure is compared to the ideal case, where sequences from a single organism (generated *in silico*) are aligned against the database in an identical manner. After that normalization factor is applied, the resulting score indicates whether the source sample contained any of the organisms in the reference database. The analysis is conducted independently on both the species and the strain level, so that if the true strain is not present in the database, the species of origin will still be identified. While many methods consider the complete taxonomic tree and assign reads to the least common ancestor, SIANN considers only two taxonomic levels: species and strain, throwing out anything that is not unique at one of those levels and thus obviating many of the confounding factors introduced by manually curated taxonomies.

The example shown in Figure 1b indicates that species-specific reads are identified as reads that align to one species (Species I, in that case) but not the other. If Species II were not present in the example shown in Figure 1b, a much larger number of reads would be assigned as “species-specific,” when in fact those regions are shared with other species. Therefore, the ability of this method to identify strain- and species-specific sequences is a direct function of the inclusion of near neighbors in the reference database. This characteristic is shared among many classification algorithms, but it is of particular note for this method when users have an opportunity to construct their own database. In order to detect a target species with a high degree of specificity (reducing false positives), it is necessary to include other related species in the reference database. Only by parallel alignment to those near neighbors can the redundant sequences be separated from the species-specific ones. For example, in order to detect *Bacillus anthracis* in a sample, it would be necessary to include other species of *Bacilli* in the reference database so that the presence of *B. cereus* or *B. thuringiensis* in a sample does not lead to a false call for *B. anthracis*.

The nomenclature of genus, species, and strain is potentially problematic because it does not correspond to a consistent degree of evolutionary distance or genomic distinctiveness. The ability to distinguish two organisms by any method using genomic sequence data is proportional to the amount of each genome that is shared or unique. One might assume that any two organisms of the same species will have a relatively predictable amount of shared genomic identity. However, some pairs of organisms from the same species may have less in common than other pairs of organisms from different species or even genera. This ambiguity impacts SIANN in two ways. If two organisms have very little genomic sequence to distinguish them, the sensitivity of SIANN to detect either one will diminish (the rate of false negatives will increase as the likelihood of sequencing unique regions decreases). Conversely, if an organism is extremely dissimilar to the near neighbors selected for the database, the specificity with which SIANN detects that organism will decline (the rate of false positives will increase as the number of related genomes available in the database decreases). For example, if a database contained only *E. coli* and *B.anthracis*, a sample containing *B. cereus* would be misidentified as contraining *B. anthracis.* In the intended use case, a database targeting *B. anthracis* would contain *B. cereus* and a number of other near neighbors to prevent that kind of misidentification. It would be convenient to say that an ideal database can be made by calculating the ideal genetic distance between all references and then finding an ideal set of organisms to make up that database, but the behavior of any database will be governed by the particular genomes of the organisms it encounters in the wild. Because not all organisms evolve in the same manner (differences in mutation rate, horizontal gene transfer, recombination, etc), the suitability of a database and method to detect a given organism can only be determined by thorough validation and benchmarking, as well as updating the reference database as needed. Users of SIANN may construct their own custom databases to include newly identified genomes or specific subsets of genomes that best suit their research interests.

Steps to construct a custom database:

1. Select a set of target organisms
2. Gather a set of genome sequences for those target organisms as well as a matched set of near neighbors
3. Using those reference genome sequences as an input, SIANN will:

a. Construct a reference index for alignment
b. Simulate a set of reads from each genome
c. Align each of those simulated read sets to all of the reference genomes
d. Calculate the proportion of each reference genome that is strain- or species-specific
e. [If two organisms do not have a minimal amount of unique sequences that exceeds the rate of sequencing error, SIANN asks that all but one of those organisms are removed from the database to eliminate redundancy. Note that the user can provide a single representative genome with multiple strain names so that the redundant strain names are not lost.]

The files contained within each SIANN database are a compressed genomic index and a list containing the proportion of each reference genome that was found to be strain- or species-specific during database construction.

To run SIANN:

1. Select a pre-made SIANN database and a set of sequences to be analyzed, and
2. SIANN will:

a. Align each of the reads against the reference genomes
b. Calculate the proportion of each reference genome that is strain- or species-specific within those reads
c. Compare that proportion to the simulated ideal case generated during database creation
d. Calculate the probability that the given results could be generated by random chance
e. Report the normalized proportion and non-parametric statistic of likelihood for each strain and species in the reference database. The normalized proportion of the genome covered by strain- or species- specific reads is the primary statistic reported by this tool.

### Benchmarking

The performance of SIANN (version 1.6) was tested in comparison to the following state-of-the-art metagenomic classification programs: LMAT (version 1.2), MetaPhlAn (version 1.7.7), and Kraken (version 0.9.1b). All of the programs in Table 1 were investigated for this effort, and three were chosen based on their ability to run on our high-performance computing cluster with an execution time and memory requirement that would be suitable to a clinical lab. Each program was run on a set of 600 simulated datasets generated by MetaSim (Richter, et al., 2008). Each dataset consisted of 15,000,000 reads (100bp single-ended) with Illumina-simulated error (fourth-degree polynomial) (Korbel, et al., 2009). The 600 datasets were broken into 12 sets of 50 replicates. Each of the 12 sets contained organisms at different levels of abundance as shown in Table 2. Organisms were specifically chosen in pairs so that the ability to distinguish these near neighbors could be determined. The abundances were staggered at 4-fold intervals so that a wide range could be evaluated. All known species of near neighbors for each of the 12 target organisms were included in the reference database used by SIANN for this benchmarking (“Target Pathogen Database”) and are shown in Appendix 1.

**Table 2.** The abundance of each target organism in each set of simulated datasets. Each set is indicated by the number in the top row, and was generated with 50 replicates.

| Organism | 1 | 2 | 3 | 4 | 5 | 6 | 7 | 8 | 9 | 10 | 11 | 12 |
| --- | --- | --- | --- | --- | --- | --- | --- | --- | --- | --- | --- | --- |
| *Bacillus anthracis* | 0.074% | 0.30% | 1.2% | 4.7% | 19% | 76% |  |  |  |  |  |  |
| *Bacillus cereus* |  |  |  |  |  |  | 0.074% | 0.30% | 1.2% | 4.7% | 19% | 76% |
| Hanta virus |  | 1.2% | 4.7% | 19% | 76% | 0.074% | 0.30% |  |  |  |  |  |
| Rift valley fever virus | 0.30% |  |  |  |  |  |  | 1.2% | 4.7% | 19% | 76% | 0.074% |
| *Clostridium botulinum* |  |  | 19% | 76% | 0.074% | 0.30% | 1.2% | 4.7% |  |  |  |  |
| *Clostridium difficile* | 1.2% | 4.7% |  |  |  |  |  |  | 19% | 76% | 0.074% | 0.30% |
| *Listeria fleischmann eii* |  |  |  | 0.074% | 0.30% | 1.2% | 4.7% | 19% | 76% |  |  |  |
| *Listeria monocytogenes* | 4.7% | 19% | 76% |  |  |  |  |  |  | 0.074% | 0.30% | 1.2% |
| Monkeypox virus |  |  |  |  | 1.2% | 4.7% | 19% | 76% | 0.074% | 0.30% |  |  |
| *Vaccinia virus* | 19% | 76% | 0.074% | 0.30% |  |  |  |  |  |  | 1.2% | 4.7% |
| *Yersinia enterocolitica* |  |  |  |  |  | 19% | 76% | 0.074% | 0.30% | 1.2% | 4.7% |  |
| *Yersinia pestis* | 76% | 0.074% | 0.30% | 1.2% | 4.7% |  |  |  |  |  |  | 19% |

Each program outputs a distinct measure. Kraken and LMAT both count the reads assigned to each taxon, MetaPhlAn calculates the abundance, and SIANN outputs a measure of the proportion of diagnostic genomic regions present. To put these measures on an even footing, we empirically calculated the false positive rate for each method over all 600 samples, at each possible measure of output. Because each dataset is made up of known organisms, any result can be classified as true or false. Therefore, for any possible result (say, 513 reads classified by LMAT or 1.6% abundance assigned by MetaPhlAn), one can calculate the proportion of calls with at least the same amount of support that were correct (True Positives/[True Positives + False Positives]), over all of the 600 datasets. That measure is commonly given as Positive Predictive Value (PPV). For each program, the results can be translated from the raw value into a PPV that is based on this empirical measure of error. The key item of interest is the PPV value for the results that we know to be true positives, the defined spike organisms. Another way of describing this approach is to say that the results of each program have been normalized to the false positive error rate that was empirically observed. If another set of samples were generated, the PPV vs. raw value curve (Figure 2) would likely fall differently, but in this case it gives us a means of comparing a diverse set of methods against the same ground truth. If method 1 detects an organism with a higher PPV than method 2 does, it means that method 1 has fewer false positives in the range that it reports true positives, which is the definition of utility in this setting.

**Figure 2.**
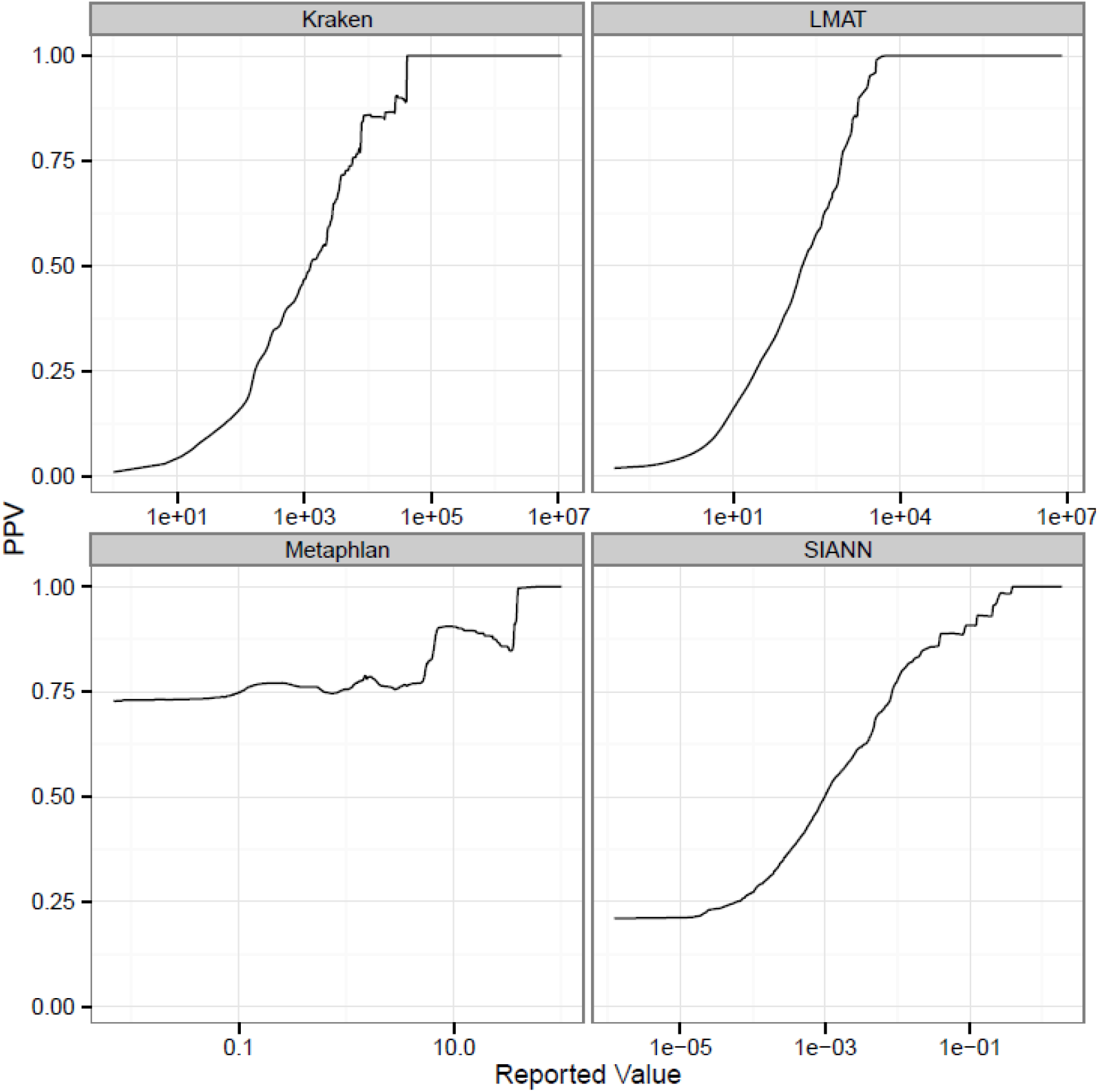
Relationship of reported value for each program (horizontal axis, log scale) to the empirically-determined Positive Predictive Value (PPV), shown on the vertical axis. While the exact values depend on the test data used, the general values at significant cutoff values (0.8, 0.9, 0.95 PPV) remain relatively constant across different datasets (data not shown).

For each method, PPV was calculated as a function of raw output value. Briefly, this was done by compiling the output for all 600 samples, labeling each result as false or true based on the sample set that it came from, and then calculating (at each possible value of output) what the proportion of TP/[TP + FP] was for results with at least that level of raw output. Some simplification steps were taken, such as focusing on the species-level assignments (for comparison with methods that do not perform strain assignment), and only taking the top hit for each species from each dataset. Custom R and BASH scripts were used for the data compilation and analysis.

## Results

The relationship of raw output value to PPV is shown for each of the four methods in Figure 2. The point at which PPV is very close to 1 (where 95% of results are true positives) is ∼41,000 reads for Kraken, ∼2,800 reads for LMAT, ∼38% abundance for MetaPhlAn, and 0.21 for SIANN. For SIANN this means that having 38% of the species-unique genome covered by reads resulted in the vast majority of calls being accurate.

For read-assignment methods (such as LMAT and Kraken), manual inspection of the results may yield a different understanding of confidence than is presented here, or in any automated analysis. For example, while each read that is assigned by LMAT and Kraken fall above a certain cutoff for species-specificity, some individual reads may be much more specific than others. One could identify a read that aligns to a single species of bacteria with 100% accuracy over its 300bp length, with the next closest match being only 90% similar. It is extremely unlikely that a 300bp exact match would arise due to random chance, and so the user could say with confidence that the organism of interest is found within the sequence data (not considering contamination, horizontal gene transfer, etc). However, such an approach is not currently implemented in an automated method, and many of the steps needed to make that assertion are performed manually by a domain expert, including alignment to near neighbors and ensuring that the read does not fall within a transposon, plasmid, etc. Therefore, while one could say that a single read is all that is needed to state with high PPV that an organism is present, the amount of reads assigned in an automated manner needed to achieve that level of PPV will number in the thousands (Fig 2).

The next phase of benchmarking was to determine how many raw input reads were needed to achieve the threshold for high PPV. To demonstrate this we plotted the known abundance of each spike organism against the PPV value generated by each method (Figure 3). Each point (an organism at a known level of abundance) is comprised of a maximum of 50 replicates, where the diameter of each point increases with an increasing number of replicates. For demonstration purposes we are showing two pairs of bacteria and three viruses. Recall that for each of the pairs of bacteria (and the two poxviruses) any sample containing one did not contain the other (as shown in Table 1). The empty boxes result from the organisms not being called at any abundance. For MetaPhlAn, that is a result of no viruses being included in the version of the reference database available for this analysis. Kraken assigned no reads to Hanta virus because viral RNA genomes were not included in this version of the reference database (personal communication with D. Wood). This emphasizes the point that a) the ability to create custom databases targeting organisms of interest can be valuable, and b) the performance of any method must be benchmarked against each potential target of interest.

**Figure 3.**
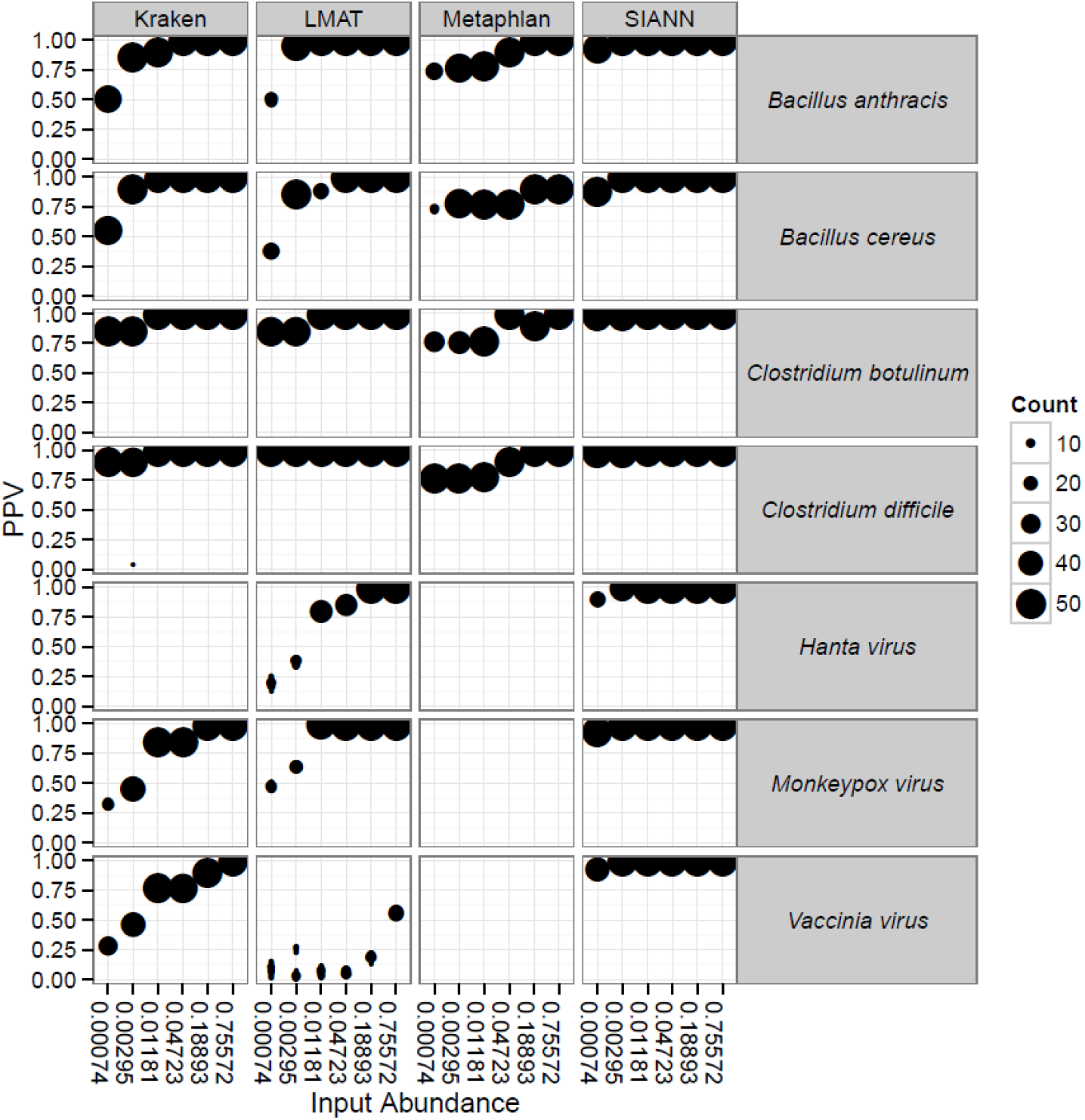
The Positive Predictive Value (PPV, vertical axis) is shown for each organism (boxes on right), at each level of known abundance (horizontal axis, see Table 2), and each program (boxes at top), across a maximum of 50 replicates (indicated by the size of each point). Note that the reference database for MetaPhlAn does not include viruses, and the reference database for Kraken does not include RNA viruses

All methods were able identify the bulk of organisms in their databases at high abundances (75% and 18%, Figure 3), however performance varied considerably at lower abundances and depended on the particular organism and method used. SIANN detected each organism at high confidence, even at levels as low as 0.3% and 0.07% of the total.

## Discussion

The process of detecting trace amounts of a specific organism in a complex mixture of DNA is challenging enough for an expert, but that pales in comparison to the difficulty of accomplishing the same certainty of detection in an automated manner. The results presented here show that SIANN rapidly detects the presence of a given set of organisms with a high degree of specificity and sensitivity. For example, at the 95% confidence (PPV) cutoff of 0.2, SIANN reliably detects all of the organisms tested here at as low as 0.3% abundance. This strong performance is likely due to the fact that SIANN is able to use a method (read alignment to whole genomes) that would be far too computationally costly if it were applied to the entire collection of known genomes. By focusing on a set of (customizable) target organisms and their near neighbors, SIANN can operate quickly and with low computational requirements while delivering highly accurate results.

SIANN is available on Illumina’s BaseSpace (www.basespace.illumina.com) as a NativeApp, with the database tested here (Appendix 1), as well as a database made from the NCBI representative set of prokaryotic genomes (ftp://ftp.ncbi.nlm.nih.gov/genomes/genome_reports/) (Appendix 2) and the complete set of NCBI viral genomes (ftp://ftp.ncbi.nlm.nih.gov/refseq/release/viral/) (Appendix 3).

BaseSpace was chosen as an appropriate release platform because while the entire set of software and dependencies can be deployed by the user from within a graphical user interface, the actual computation takes place in a controlled ‘cloud’ environment. Such a distribution strategy obviates the need to satisfy the multiple software or OS dependencies that often arises with academic computational methods. Results for SIANN are compiled into a report format, showing both the organisms that surpass 95% confidence, as well as the closest strain match for each species. The default view masks the raw data output, so that the results are human-readable and do not present extraneous information. While the code for execution and database-construction on a users system is available from Signature Science, LLC, additional databases on the BaseSpace platform can be made available upon request.

There is a neverending list of questions that one could ask of metagenomic sequencing data generated from important samples. Instead of answering them all, we demonstrate a technique with a very narrow focus that is able to report with a high degree of confidence whether a given set of organisms is present in a sample. These results are presented to the user in a comprehensible format, and accessible on a commonly-used web platform. The world of bioinformatics will continue to progress and develop more sophisticated tools for metagenomic analysis, and we hope that the utility of SIANN will convince others to package and benchmark their tools in a way that they can be used with confidence by the larger public, as well as the research community.

## Appendices

## Appendix 1: Target Pathogen Database

Arenaviridae Arenavirus Golden-Gate-virus

Arenaviridae Arenavirus Lujo-virus

Arenaviridae Flexal-virus segment-L

Arenaviridae New-world-arenaviruses Allpahuayo-virus

Arenaviridae New-world-arenaviruses Chapare-virus

Arenaviridae New-world-arenaviruses Guanarito-virus

Arenaviridae New-world-arenaviruses Junin-virus

Arenaviridae New-world-arenaviruses Machupo-virus

Arenaviridae New-world-arenaviruses Sabia-virus

Arenaviridae New-world-arenaviruses Tacaribe-virus

Arenaviridae New-world-arenaviruses Whitewater-Arroyo-virus

Arenaviridae Old-world-arenaviruses Ippy-virus

Arenaviridae Old-world-arenaviruses Lassa-virus

Arenaviridae Old-world-arenaviruses Mopeia-virus-AN20410

Asfarviridae African-swine-fever-virus Benin-971-pathogenic-isolate

Asfarviridae African-Swine-Fever-Virus

Bacillus anthracis A2012 Bant 02 1

Bacillus anthracis Ames Ancestor

Bacillus anthracis Sterne

Bacillus cereus 03BB102

Bacillus cereus AH187

Bacillus cereus AH820

Bacillus cereus ATCC 14579

Bacillus cereus B4264

Bacillus cereus F65185

Bacillus cytotoxicus NVH 391-98

Bacillus mycoides DSM 2048

Bacillus mycoides Rock1-4

Bacillus thuringiensis BMB171

Bacillus thuringiensis Bt407

Bacillus thuringiensis HD-771

Bacillus thuringiensis serovar chinensis CT-43

Bacillus thuringiensis serovar konkukian 97-27

Brucella abortus A13334

Brucella ceti B1 94

Brucella ceti M13 05 1 supercont1 22

Brucella melitensis ATCC 23457

Brucella melitensis bv 1 str 16M

Brucella ovis ATCC 25840

Brucella suis 1330

Brucella suis ATCC 23445

Bunyaviridae Akabane-virus segment-M

Bunyaviridae Hantavirus Andes-virus

Bunyaviridae Hantavirus Dobrava-Belgrade-virus-strain-DOBV-Ano-Poroia-Afl9-1999

Bunyaviridae Hantavirus Hantaan-virus

Bunyaviridae Hantavirus Puumala-virus

Bunyaviridae Hantavirus Seoul-virus-strain-Seoul-80-39-clone-1

Bunyaviridae Hantavirus Sin-Nombre-virus

Bunyaviridae Hantavirus Thottapalayam-virus

Bunyaviridae Hantavirus Tula-virus

Bunyaviridae Nairovirus Crimean-Congo-hemorrhagic-fever-virus

Bunyaviridae Nairovirus Dugbe-virus

Bunyaviridae Phlebovirus Rift-Valley-fever-virus

Burkholderia cenocepacia HI2424

Burkholderia cenocepacia J2315

Burkholderia cenocepacia MC0-3

Burkholderia cepacia GG4

Burkholderia gladioli BSR3

Burkholderia glumae BGR1

Burkholderia mallei 2002721280

Burkholderia mallei ATCC 10399

Burkholderia mallei NCTC 10247

Burkholderia mallei SAVP1

Burkholderia multivorans ATCC 17616

Burkholderia oklahomensis C6786

Burkholderia oklahomensis EO147

Burkholderia pseudomallei 1026b

Burkholderia pseudomallei 1106a

Burkholderia pseudomallei 1710b

Burkholderia pseudomallei 668

Burkholderia pseudomallei BPC006

Burkholderia pseudomallei K96243

Burkholderia pyrrocinia CH-67

Burkholderia thailandensis ATCC 700388

Burkholderia thailandensis E264

Burkholderia thailandensis MSMB121

Campylobacter coli JV20 Campylobacter fetus subsp fetus 82-40

Campylobacter jejuni RM1221

Campylobacter jejuni subsp doylei 26997

Campylobacter jejuni subsp jejuni 81-176

Campylobacter jejuni subsp jejuni NCTC 11168 ATCC 700819

Campylobacter upsaliensis JV21

Clostridium acetobutylicum DSM 1731

Clostridium botulinum A str ATCC 3502

Clostridium botulinum BKT015925

Clostridium botulinum B str Eklund 17B

Clostridium botulinum E3 str Alaska E43

Clostridium botulinum F str 230613

Clostridium botulinum H04402 065

Clostridium difficile 2007855

Clostridium difficile 630

Clostridium difficile BI1

Clostridium difficile BI9

Clostridium perfringens ATCC 13124

Clostridium perfringens SM101

Clostridium perfringens str 13

Clostridium symbiosum WAL-14163

Clostridium symbiosum WAL-14673

Clostridium tetani E88

Clostridium thermocellum ATCC 27405

Clostridium thermocellum DSM 1313

Clostridium tunisiense TJ C661

Clostridium ultunense Esp

Coccidioides immitis H5384

Coccidioides immitis RMSCC 2394

Coccidioides immitis RS

Coccidioides posadasii C735 delta SOWgp

Coccidioides posadasii CPA 0001

Coccidioides posadasii CPA 0020

Coronaviridae Alphacoronavirus Bat-coronavirus-HKU2

Coronaviridae Alphacoronavirus Feline-infectious-peritonitis-virus

Coronaviridae Alphacoronavirus Human-coronavirus-229E

Coronaviridae Alphacoronavirus TGEV-Purdue-P115

Coronaviridae Bafinivirus White-bream-virus

Coronaviridae Betacoronavirus Bovine-coronavirus

Coronaviridae Betacoronavirus Murine-hepatitis-virus-strain-A59

Coronaviridae Betacoronavirus Murine-hepatitis-virus-strain-JHM

Coronaviridae Betacoronavirus SARS-coronavirus

Coronaviridae Coronavirinae Munia-coronavirus-HKU13-3514

Coronaviridae Gammacoronavirus Avian-infectious-bronchitis-virus

Coronaviridae Gammacoronavirus Turkey-coronavirus

Coronaviridae Torovirus Breda-virus

Coronaviridae unclassified-coronaviruses Bat-coronavirus-BM48-31-BGR-2008

Coronaviridae unclassified-coronaviruses Bovine-respiratory-coronavirus-AH187

Coronaviridae unclassified-coronaviruses Human-enteric-coronavirus-strain-4408

Coxiella burnetii CbuG Q212

Coxiella burnetii Dugway 5J108 111

Coxiella burnetii RSA 493 Diplorickettsia massiliensis 20B CS 1

Ehrlichia canis str Jake Ehrlichia chaffeensis str Arkansas

Ehrlichia chaffeensis str Sapulpa ctg90

Ehrlichia ruminantium str Gardel

Ehrlichia ruminantium str Welgevonden

Escherichia coli APEC O1

Escherichia coli BL21 DE3

Escherichia coli B str REL606

Escherichia coli ETEC H10407

Escherichia coli O104H4 str 2011C-3493

Escherichia coli O157H7 str EC4115

Escherichia coli O157H7 str Sakai

Escherichia coli O7K1 str CE10

Escherichia coli O83H1 str NRG 857C

Escherichia coli str K-12 substr MG1655

Filoviridae Ebolavirus Bundibugyo-ebolavirus

Filoviridae Ebolavirus Cote-dIvoire-ebolavirus

Filoviridae Ebolavirus Ebola-virus-Mayinga-Zaire

Filoviridae Ebolavirus Reston-ebolavirus

Filoviridae Ebolavirus Sudan-ebolavirus

Filoviridae Marburgvirus Lake-Victoria-marburgvirus-Musoke

Flaviviridae Alkhurma-virus

Flaviviridae Classical-swine-fever-virus

Flaviviridae Dengue-virus 1

Flaviviridae Dengue-virus 2

Flaviviridae Dengue-virus 3

Flaviviridae Dengue-virus 4

Flaviviridae Japanese-encephalitis-virus genome

Flaviviridae Karshi-virus

Flaviviridae Langat-virus

Flaviviridae Louping-ill-virus

Flaviviridae Murray-Valley-encephalitis-virus

Flaviviridae Omsk-hemorrhagic-fever-virus

Flaviviridae Powassan-virus

Flaviviridae St-Louis-encephalitis-virus

Flaviviridae Tick-borne-encephalitis-virus

Flaviviridae Usutu-virus

Flaviviridae West-Nile-virus

Flaviviridae Yellow-fever-virus

Francisella cf novicida Fx1

Francisella noatunensis subsp orientalis str Toba 04

Francisella novicida U112

Francisella philomiragia subsp philomiragia ATCC 25015

Francisella philomiragia subsp philomiragia ATCC 25017

Francisella tularensis subsp holarctica F92

Francisella tularensis subsp mediasiatica FSC147

Francisella tularensis subsp tularensis FSC198

Herpesviridae Alcelaphine-herpesvirus 1

Herpesviridae Macacine-herpesvirus 1

Listeria fleischmannii LU2006-1 c88

Listeria innocua Clip11262

Listeria innocua FSL J1-023

Listeria ivanovii FSL F6-596

Listeria ivanovii subsp ivanovii PAM 55

Listeria marthii FSL S4-120

Listeria monocytogenes 07PF0776

Listeria monocytogenes 08-5578

Listeria monocytogenes 10403S

Listeria monocytogenes ATCC 19117

Listeria monocytogenes Finland 1998

Listeria monocytogenes FSL R2-561

Listeria monocytogenes La111

Listeria seeligeri FSL N1-067

Listeria seeligeri serovar 12b str SLCC3954

Listeria welshimeri serovar 6b str SLCC5334

Paramyxoviridae Avulavirus Newcastle-disease-virus-B1

Paramyxoviridae Henipavirus Hendra-virus

Paramyxoviridae Henipavirus Nipah-virus

Paramyxoviridae Menangle-virus

Paramyxoviridae Morbillivirus Measles-virus

Paramyxoviridae Peste-des-petits-ruminants-virus

Paramyxoviridae Respirovirus Human-parainfluenza-virus-1

Paramyxoviridae Respirovirus Human-parainfluenza-virus-3

Paramyxoviridae Respirovirus Sendai-virus

Paramyxoviridae Rinderpest-virus strain-Kabete-O

Paramyxoviridae Rubulavirus Human-parainfluenza-virus-2

Paramyxoviridae Rubulavirus Human-parainfluenza-virus-4a

Paramyxoviridae Rubulavirus Mumps-virus

Picornaviridae Foot-and-mouth-disease-virus -type-O

Picornaviridae Swine-vesicular-disease-virus strain-HK70

Picornaviridae Swine-vesicular-disease-virus strain-NET192

Poxviridae Avipoxvirus Fowlpox-virus

Poxviridae Crocodylidpoxvirus Nile-crocodilepox-virus

Poxviridae Goatpox-virus Pellor

Poxviridae Leporipoxvirus Myxoma-virus

Poxviridae Lumpy-skin-disease-virus NI-2490

Poxviridae Molluscipoxvirus Molluscum-contagiosum-virus-subtype-1

Poxviridae Orthopoxvirus Camelpox-virus

Poxviridae Orthopoxvirus Cowpox-virus

Poxviridae Orthopoxvirus Ectromelia-virus

Poxviridae Orthopoxvirus Monkeypox-virus-Zaire-96-I-16

Poxviridae Orthopoxvirus Taterapox-virus

Poxviridae Orthopoxvirus Vaccinia-virus

Poxviridae Orthopoxvirus Variola-virus

Poxviridae Sheeppox-virus 17077-99

Poxviridae Suipoxvirus Swinepox-virus

Puccinia graminis f sp tritici CRL 75-36-700-3

Ralstonia pickettii 12D

Ralstonia pickettii 12J

Ralstonia solanacearum CFBP2957

Ralstonia solanacearum CMR15

Ralstonia solanacearum GMI1000

Rathayibacter toxicus DSM 7488

Reoviridae African-horsesickness-virus segment-10

Rhabdoviridae Vesicular-stomatitis-Indiana-virus

Rhabdoviridae Vesicular-stomatitis-virus strain-NJ2075212NM

Rickettsia bellii OSU 85-389

Rickettsia conorii str Malish 7

Rickettsia prowazekii str Breinl

Rickettsia prowazekii str RpGvF24

Rickettsia rickettsii str Arizona

Rickettsia typhi str B9991CWPP

Rickettsiella grylli gcontig 634

Salmonella bongori N268-08

Salmonella bongori NCTC 12419

Salmonella enterica subsp arizonae serovar 62z4z23-str RSK2980

Salmonella enterica subsp enterica serovar Dublin str CT 02021853

Salmonella enterica subsp enterica serovar Newport str SL254

Salmonella enterica subsp enterica serovar Paratyphi A str ATCC 9150

Salmonella enterica subsp enterica serovar Typhimurium str LT2

Salmonella enterica subsp enterica serovar Typhi str Ty2

Shigella boydii CDC 3083-94

Shigella boydii Sb227

Shigella dysenteriae Sd197

Shigella flexneri 2002017

Shigella flexneri 2a str 2457T

Shigella flexneri 2a str 301

Shigella flexneri 5 str 8401

Shigella sonnei 53G

Shigella sonnei Ss046

Staphylococcus arlettae CVD059 SARL c230

Staphylococcus aureus 04-02981

Staphylococcus aureus 08BA02176

Staphylococcus aureus subsp aureus N315

Staphylococcus aureus subsp aureus NCTC 8325

Staphylococcus aureus subsp aureus TW20

Staphylococcus capitis QN1 Contig63

Staphylococcus capitis SK14

Staphylococcus caprae C87

Staphylococcus carnosus subsp carnosus TM300

Staphylococcus epidermidis ATCC 12228

Staphylococcus epidermidis RP62A

Staphylococcus equorum subsp equorum Mu2

Staphylococcus haemolyticus JCSC1435

Staphylococcus hominis SK119

Staphylococcus hominis subsp hominis C80

Staphylococcus lugdunensis HKU09-01

Staphylococcus lugdunensis N920143

Togaviridae Alphavirus Barmah-Forest-virus Togaviridae Chikungunya-virus

Togaviridae EEEV-complex Eastern-equine-encephalitis-virus

Togaviridae Rubivirus Rubella-virus

Togaviridae SFV-complex O-nyong-nyong-virus

Togaviridae Venezuelan-equine-encephalitis-virus

Togaviridae WEEV-complex Sindbis-virus

Togaviridae Western-equine-encephalomyelitis-virus

Xanthomonas albilineans GPE PC73

Xanthomonas axonopodis Xac29-1

Xanthomonas oryzae pv oryzae KACC 10331

Xanthomonas oryzae pv oryzae MAFF 311018

Xanthomonas oryzae pv oryzae PXO99A

Xanthomonas vasicola pv vasculorum NCPPB 1326 scf 9767 4580

Yersinia aldovae ATCC 35236

Yersinia bercovieri ATCC 43970

Yersinia enterocolitica IP 10393

Yersinia enterocolitica IP2222

Yersinia enterocolitica subsp enterocolitica 8081

Yersinia enterocolitica subsp palearctica 105 5R

Yersinia frederiksenii ATCC 33641

Yersinia intermedia ATCC 29909

Yersinia kristensenii ATCC 33638

Yersinia mollaretii ATCC 43969

Yersinia pestis A1122

Yersinia pestis Antiqua

Yersinia pestis KIM 10

Yersinia pestis Pestoides F

Yersinia pseudotuberculosis IP 31758

Yersinia pseudotuberculosis IP 32953

Yersinia pseudotuberculosis PB1

Yersinia pseudotuberculosis YPIII

Yersinia rohdei ATCC 43380

Yersinia ruckeri ATCC 29473

## Appendix 2: Viral Database

Abaca bunchy top virus DNA-C

Abaca bunchy top virus DNA-M

Abaca bunchy top virus DNA-N

Abaca bunchy top virus DNA-R

Abaca bunchy top virus DNA-S

Abaca bunchy top virus segment 2

Abalone shriveling syndrome-associated virus

Abutilon Brazil virus DNA A

Abutilon Brazil virus DNA B

Abutilon mosaic virus DNA A

Abutilon mosaic virus DNA B

Acanthocystis turfacea Chlorella virus 1

Acheta domesticus densovirus

Acholeplasma phage L2

Acholeplasma phage MV-L1

Acidianus bottle-shaped virus

Acidianus filamentous virus 1

Acidianus filamentous virus 2

Acidianus filamentous virus 3

Acidianus filamentous virus 6

Acidianus filamentous virus 7

Acidianus filamentous virus 8

Acidianus filamentous virus 9

Acidianus rod-shaped virus 1

Acidianus spindle-shaped virus 1

Acidianus two-tailed virus

Actinomyces phage Av-1

Actinoplanes phage phiAsp2

Acyrthosiphon pisum bacteriophage APSE-1

Adeno-associated virus-1

Adeno-associated virus-2

Adeno-associated virus-3

Adeno-associated virus-4

Adeno-associated virus 5

Adeno-associated virus-7

Adeno-associated virus-8

Adoxophyes honmai NPV

Adoxophyes orana granulovirus

Adoxophyes orana nucleopolyhedrovirus

Aedes aegypti densovirus

Aedes albopictus densovirus

Aedes taeniorhynchus iridescent virus

Aeromonas phage 25

Aeromonas phage 31

Aeromonas phage 44RR2.8t

Aeromonas phage Aeh1

Aeromonas phage phiO18P african cassava mosaic virus DNA A african cassava mosaic virus DNA B

African green monkey polyomavirus

African swine fever virus

Ageratum enation virus

Ageratum leaf Cameroon betasatellite

Ageratum leaf curl virus-G52

Ageratum yellow vein China virus-associated DNA beta

Ageratum yellow vein Chinavirus

Ageratum yellow vein Hualian virus-TaiwanHsinchutom2003 DNA A

Ageratum yellow vein Sri Lanka virus segment A

Ageratum yellow vein Taiwan virus

Ageratum yellow vein virusassociated DNA beta

Ageratum yellow veinvirus

Agrotis ipsilon multiple nucleopolyhedrovirus

Agrotis segetum granulovirus

Agrotis segetum nucleopolyhedrovirus

Alcelaphine herpesvirus 1

Aleutian mink disease virus

Allamanda leaf curl virus DNA-A

Alternanthera yellow vein virus DNA-A

Alternanthera yellow vein virus satellite DNA beta

Ambystoma tigrinum virus

Amsacta moorei entomopoxvirus L

Anguillid herpesvirus 1

Anopheles gambiae densonucleosis virus

Antheraea pernyi nucleopolyhedrovirus

Anticarsia gemmatalis nucleopolyhedrovirus

Archaeal BJ1 virus

Ateline herpesvirus 3

Autographa californica nucleopolyhedrovirus

Avian adeno-associated virus

ATCC VR-865

Avian adeno-associated virus strain DA-1

Avian endogenous retrovirus EAV-HP

Azospirillum phage Cd

Bacillus phage 0305phi8-36

Bacillus phage AP50

Bacillus phage B103

Bacillus phage Bam35c

Bacillus phage BCJA1c

Bacillus phage Cherry

Bacillus phage Fah

Bacillus phage GA-1

Bacillus phage Gamma

Bacillus phage GIL16c

Bacillus phage IEBH

Bacillus phage phi105

Bacillus phage phi29

Bacillus phage SPBc2

Bacillus phage SPO1

Bacillus phage SPP1

Bacillus phage TP21-L

Bacillus phage WBeta

Bacillus prophage phBC6A51

Bacillus prophage phBC6A52

Bacillus virus 1

Bacteriophage Aaphi23

Bacteriophage APSE-2

Bacteriophage PSA

Bacteriophage RB32

Bacteroides phage B40-8

Banana bunchy top virus DNA C

Banana bunchy top virus DNA M

Banana bunchy top virus DNA N

Banana bunchy top virus DNA R

Banana bunchy top virus DNA S

Banana bunchy top virus DNA U3

Banana streak GF virus

Banana streak Mysore virus

Banana streak OL virus

Banana streak virus genome

Banana streak virus strain Acuminata Vietnam

Bandicoot papillomatosis carcinomatosis virus type 1

Bandicoot papillomatosis carcinomatosis virus type 2

Bat adeno-associated virus YNM

Bdellovibrio phage phiMH2K

Beak and feather disease virus

Bean calico mosaic virus DNA A

Bean calico mosaic virus DNA B

Bean dwarf mosaic virus DNA A

Bean dwarf mosaic virus DNA B

Bean golden mosaic virus DNA A

Bean golden mosaic virus DNA B

Bean golden yellow mosaic virus DNA A

Bean golden yellow mosaic virus DNA B

Bean yellow dwarf virus putative genes V1

Beet curly top Iran virus-K

Beet curly top virus-California Logan

Beet mild curly top virus-Worland4

Beet severe curly top virus-Cfh

Begomovirus-associated DNA II

Begomovirus-associated DNA-III

Bettongia penicillata papillomavirus 1

Bhendi yellow vein

Bhubhaneswar virus DNA-A

Bhendi yellow vein Delhi virus 2004New Delhi DNA-A

Bhendi yellow vein mosaic virus-associated DNA beta

Bhendi yellow vein mosaic Virus

Bitter gourd leaf curl disease-associated DNA beta

BK polyomavirus

Blainvillea yellow spot virus DNA-A

Blainvillea yellow spot virus DNA-B

Blattella germanica densovirus

Blueberry red ringspot virus

Bocavirus gorillaGBoV12009

Bombyx mandarina nucleopolyhedrovirus

Bombyx mori densovirus 5

Bombyx mori NPV

Bordetella phage BIP-1

Bordetella phage BMP-1

Bordetella phage BPP-1

Bougainvillea spectabilis chlorotic vein-banding virus

Bovine adeno-associated virus

Bovine adenovirus A

Bovine adenovirus B

Bovine adenovirus D

Bovine ephemeral fever virus

Bovine foamy virus

Bovine herpesvirus 1

Bovine herpesvirus 4 long unique region

Bovine herpesvirus 5

Bovine papillomavirus-1

Bovine papillomavirus 3

Bovine papular stomatitis virus

Bovine parvovirus 2

Bovine Parvovirus

Bovine polyomavirus

Burkholderia ambifaria phage BcepF1

Burkholderia phage Bcep 176

Burkholderia phage Bcep1

Burkholderia phage Bcep22

Burkholderia phage Bcep43

Burkholderia phage Bcep781

Burkholderia phage BcepB1A

Burkholderia phage BcepC6B

Burkholderia phage BcepGomr

Burkholderia phage BcepIL02

Burkholderia phage BcepMu

Burkholderia phage BcepNazgul

Burkholderia phage BcepNY3

Burkholderia phage KS10

Burkholderia phage KS9

Burkholderia phage phi1026b

Burkholderia phage phi644-2 chromosome

Burkholderia phage phiE12-2 chromosome

Burkholderia phage phiE125

Burkholderia phage phiE202 chromosome

Burkholderia phage phiE255 chromosome

Burkholderia prophage phi52237

Cabbage leaf curl virus DNA A

Cabbage leaf curl virus DNA B

Cacao swollen shoot virus

California sea lion anellovirus

California sea lion polyomavirus 1

Callitrichine herpesvirus 3 strain CJ0149

Camelpox virus

Campoletis sonorensis ichnovirus chromosome segment W

Campoletis sonorensis ichnovirus segment B

Campoletis sonorensis ichnovirus segment C

Campoletis sonorensis ichnovirus segment D

Campoletis sonorensis ichnovirus segment E

Campoletis sonorensis ichnovirus segment F

Campoletis sonorensis ichnovirus segmentG2

Campoletis sonorensis ichnovirus segment G

Campoletis sonorensis ichnovirus segment H

Campoletis sonorensis ichnovirus segmentI2

Campoletis sonorensis ichnovirus segment I

Campoletis sonorensis ichnovirus segment J

Campoletis sonorensis ichnovirus segment L

Campoletis sonorensis ichnovirus segment M

Campoletis sonorensis ichnovirus segment N

Campoletis sonorensis ichnovirus segment O1

Campoletis sonorensis ichnovirus segment P

Campoletis sonorensis ichnovirus segment Q

Campoletis sonorensis ichnovirus segment T

Campoletis sonorensis ichnovirus segment U

Campoletis sonorensis ichnovirus segment V

Campoletis sonorensis ichnovirus segment Z

Campoletis sonorensis ichnovirus superhelical segment A

Campoletis sonorensis ichnovirus superhelical segment aprime

Canary circovirus

Canarypox virus

Canine adenovirus Canine minute virus

Canine oral papillomavirus

Canine papillomavirus 2

Canine papillomavirus 3

Canine papillomavirus 4

Canine papillomavirus 6

Canine parvovirus

Capra hircus papillomavirus type 1

Cardiospermum yellow leaf curl virus satellite DNA beta

Caretta caretta papillomavirus 1

Carnation etched ring virus

Casphalia extranea densovirus

Cassava vein mosaic virus

Cauliflower mosaic virus

Cercopithecine herpesvirus 2

Cercopithecine herpesvirus 5 strain 2715

Cercopithecine herpesvirus 9

Cestrum yellow leaf curling virus

Chaetoceros salsugineum DNA virus

Chayote yellow mosaic virus

Chicken anemia virus

Chickpea chlorotic dwarf virus

Chilli leaf curl disease associated sequence virion

Chilli leaf curl Multan alphasatellite

Chilli leaf curl virus

Chino del tomate virus DNA A

Chino del tomate virus DNA B

Chlamydia phage 3

Chlamydia phage 4

Chlamydia phage Chp1

Chlamydia phage Chp2

Chlamydia phage PhiCPG1

Chloris striate mosaic virus

Choristoneura fumiferana DEF MNPV

Choristoneura fumiferana MNPV

Choristoneura occidentalis granulovirus

Chrysodeixis chalcites nucleopolyhedrovirus

Circovirus-like genome BBC-A

Circovirus-like genome CB-A

Circovirus-like genome CB-B

Circovirus-like genome RW-A

Circovirus-like genome RW-B

Circovirus-like genome RW-C

Circovirus-like genome RW-D

Circovirus-like genome RW-E

Circovirus-like genome SAR-A

Circovirus-like genome SAR-B

Citrus psorosis virus RNA1

Citrus psorosis virus RNA2

Citrus psorosis virus RNA3

Citrus yellow mosaic virus

Clanis bilineata nucleopolyhedrosis virus

Clavibacter phage CMP1

Clerodendron yellow mosaic virus

Clerodendrum golden mosaic

China virus DNA A

Clerodendrum golden mosaic

China virus DNA B

Clerodendrum golden mosaic virus DNA-A

Clerodendrum golden mosaic virus DNA-B

Clostridium phage 39-O

Clostridium phage c-st

Clostridium phage phi3626

Clostridium phage phiC2

Clostridium phage phi CD119

Clostridium phage phiCD27

Clostridium phage phiCTP1

Coconut foliar decay virus

Columbid circovirus

Commelina yellow mottle virus

Corchorus golden mosaic virus DNA-A

Corchorus golden mosaic virus DNA-B

Corchorus yellow spot virus DNA A

Corchorus yellow spot virus DNA B

Corchorus yellow vein virus-Hoa Binh DNA A

Corchorus yellow vein virus-Hoa Binh DNA B

Corynebacterium phage BFK20

Corynebacterium phage P1201

Cotesia congregata bracovirus segment Circle10

Cotesia congregata bracovirus segment Circle11

Cotesia congregata bracovirus segment Circle12

Cotesia congregata bracovirus segment Circle13

Cotesia congregata bracovirus segment Circle14

Cotesia congregata bracovirus segment Circle15

Cotesia congregata bracovirus segment Circle17

Cotesia congregata bracovirus segment Circle19

Cotesia congregata bracovirus segment circle1

Cotesia congregata bracovirus segment Circle20

Cotesia congregata bracovirus segment Circle21

Cotesia congregata bracovirus segment Circle22

Cotesia congregata bracovirus segment Circle23

Cotesia congregata bracovirus segment Circle25

Cotesia congregata bracovirus segment Circle26

Cotesia congregata bracovirus segment circle2

Cotesia congregata bracovirus segment Circle30

Cotesia congregata bracovirus segment Circle31

Cotesia congregata bracovirus segment Circle32

Cotesia congregata bracovirus segment Circle33

Cotesia congregata bracovirus segment Circle35

Cotesia congregata bracovirus segment Circle36

Cotesia congregata bracovirus segment circle3

Cotesia congregata bracovirus segment Circle4

Cotesia congregata bracovirus segment Circle5

Cotesia congregata bracovirus segment Circle9

Cotesia congregata virus segment Circle18

Cotesia congregata virus segment Circle6

Cotesia congregata virus segment Circle7

Cotesia congregata virus segment Circle8

Cotton leaf crumple geminivirus DNA B

Cotton leaf crumple virus DNA A

Cotton leaf curl Alabad virus

Cotton leaf curl Bangalore virus-associated DNA beta

Cotton leaf curl Bangalore virus segment A

Cotton leaf curl Burewala alphasatellite

Cotton leaf curl Burewala betasatellite

Cotton leaf curl Burewala virus-IndiaVehari2004

Cotton leaf curl Gezira alphasatellite

Cotton leaf curl Gezira beta

Cotton leaf curl Gezira Betasatellite extrachromosomal

Cotton leaf curl Gezira virus

Cotton leaf curl Kokhran virus

Cotton leaf curl Multan Virus

Cotton leaf curl Multan virus satellite DNA beta

Cotton leaf curl Multan virus satellite U36-1

Cotton leaf curl Rajasthan virus segment A

Cotton leaf curl virus-associated DNA beta

Cottontail rabbit papillomavirus

Cowpea severe leaf curl-associated DNA beta

Cowpox virus Crassocephalum yellow vein virus-Jinghong

Croton yellow vein mosaic alphasatellite

Croton yellow vein mosaic virus

Croton yellow vein mosaic Virus satellite DNA beta

Croton yellow vein virus Crow polyomavirus

Cryptophlebia leucotreta granulovirus

Cucurbita yellow vein virus-associated DNA beta

Cucurbit leaf crumple virus DNA A

Cucurbit leaf crumple virus DNA B

Culex nigripalpus NPV

Culex pipiens densovirus Cyanophage PSS2

Cyanophage Syn5 Cycad leaf necrosis virus

Cydia pomonella granulovirus

Cyprinid herpesvirus 3

Deer papillomavirus

Deerpox virus W-1170-84

Deerpox virus W-848-83

Deftia phage phiW-14

Dendrolimus punctatus densovirus

Desmodium leaf distortion virus DNA A

Desmodium leaf distortion virus DNA B

Diadromus pulchellus ascovirus 4a

Diatraea saccharalis densovirus

Dicliptera yellow mottle virus DNA A

Dicliptera yellow mottle virus DNA B

Digitaria streak virus

Dioscorea bacilliform virus

Dolichos yellow mosaic virus

Dracaena mottle virus

Duck adenovirus A

Duck circovirus

Duck hepatitis B virus

East African cassava mosaic Cameroon virus DNA A

East African cassava mosaic Cameroon virus DNA B

East African cassava mosaic Kenya virus DNA A

East African cassava mosaic Kenya virus DNA B

East African cassava mosaic virus DNA A

East African cassava mosaic virus DNA B

East African cassava mosaic Zanzibar virus DNA-A

East African cassava mosaic Zanzibar virus DNA B

Ecotropis obliqua NPV

Ectocarpus siliculosus virus 1

Ectromelia virus

Emiliania huxleyi virus 86

Emilia yellow vein virus-associated DNA beta

Emilia yellow vein virus-Fz1

Enterobacteria phage 13a

Enterobacteria phage 933W

Enterobacteria phage alpha3

Enterobacteria phage BA14

Enterobacteria phage BP-4795

Enterobacteria phage EcoDS1

Enterobacteria phage EPS7

Enterobacteria phage epsilon15

Enterobacteria phage ES18

Enterobacteria phage Felix 01

Enterobacteria phage Fels-2

Enterobacteria phage G4

Enterobacteria phage HK022

Enterobacteria phage HK620

Enterobacteria phage HK97

Enterobacteria phage I2-2

Enterobacteria phage ID18

Enterobacteria phage ID2 MoscowID2001

Enterobacteria phage If1

Enterobacteria phage Ike

Enterobacteria phage IME08

Enterobacteria phage JK06

Enterobacteria phage JS10

Enterobacteria phage JS98

Enterobacteria phage JSE

Enterobacteria phage K1-5

Enterobacteria phage K1E

Enterobacteria phage K1F

Enterobacteria phage lambda

Enterobacteria phage LKA1

Enterobacteria phage M13

Enterobacteria phage Min27

Enterobacteria phage Mu

Enterobacteria phage N15

Enterobacteria phage N4

Enterobacteria phage P1

Enterobacteria phage P22 virus

Enterobacteria phage P2 virus

Enterobacteria phage P4

Enterobacteria phage Phi1

Enterobacteria phage phiEco32

Enterobacteria phage phiEcoM-GJ1

Enterobacteria phage phiP27

Enterobacteria phage phiX174 sensu lato

Enterobacteria phage PRD1

Enterobacteria phage PsP3

Enterobacteria phage RB14

Enterobacteria phage RB16

Enterobacteria phage RB43

Enterobacteria phage RB49

Enterobacteria phage RB51

Enterobacteria phage RB69

Enterobacteria phage RTP

Enterobacteria phage Sf6

Enterobacteria phage SfV

Enterobacteria phage SP6

Enterobacteria phage SSL-2009a

Enterobacteria phage ST104

Enterobacteria phage St-1

Enterobacteria phage ST64T

Enterobacteria phage T1

Enterobacteria phage T3

Enterobacteria phage T4

Enterobacteria phage T5

Enterobacteria phage T7

Enterobacteria phage TLS

Enterobacteria phage VT2-Sakai

Enterobacteria phage WA13

Enterobacteria phage WV8

Enterobacteria phage YYZ-2008

Enterococcus faecalis 62 chromosome

Enterococcus faecalis 62 plasmid EF62pA

Enterococcus faecalis 62 plasmid EF62pB

Enterococcus faecalis 62 plasmid EF62pC

Enterococcus phage EF62phi

Enterococcus phage EFAP-1

Enterococcus phage phiEf11

Enterococcus phage phiEF24C

Enterococcus phage phiFL1A

Enterococcus phage phiFL2A

Enterococcus phage phiFL3A

Enterococcus phage phiFL4A

Enzootic nasal tumour virus of goats

Epiphyas postvittana NPV

Equid herpesvirus 1

Equid herpesvirus 2

Equid herpesvirus 4

Equid herpesvirus 9

Equine papillomavirus 2

Eragrostis curvula streak virus

Eragrostis streak virus

Erectites yellow mosaic virus DNA-A

Erectites yellow mosaic virus satellite DNA beta

Erethizon dorsatum papillomavirus type 1

Erinaceus europaeus papillomavirus

Erwinia amylovora phage Era103

Erwinia phage phiEa21-4

Escherichia coli bacteriophage rv5

Escherichia phage D108

Escherichia phage phiV10

Eupatorium vein clearing virus

Eupatorium yellow vein associated DNA beta

Eupatorium yellow vein virus

Euphorbia leaf curl virus DNA A

Euphorbia yellow mosaic virus DNA A

Euphorbia yellow mosaic virus DNA B

Euproctis pseudoconspersa nucleopolyhedrovirus

European elk papillomavirus

Faba bean necrotic stunt virus DNA C

Faba bean necrotic stunt virus DNA M

Faba bean necrotic stunt virus DNA N

Faba bean necrotic stunt virus DNA R

Faba bean necrotic stunt virus DNA S

Faba bean necrotic stunt virus DNA U1

Faba bean necrotic stunt virus DNA U2

Faba bean necrotic stunt virus DNA U4

Faba bean necrotic yellows virus DNA 10

Faba bean necrotic yellows virus DNA-1

Faba bean necrotic yellows virus DNA 2

Faba bean necrotic yellows virus DNA 4

Faba bean necrotic yellows virus DNA 5

Faba bean necrotic yellows virus DNA 7

Faba bean necrotic yellows virus DNA 8

Faba bean necrotic yellows virus DNA 9

Faba bean necrotic yellows Virus

Feldmannia species virus Felid herpesvirus 1

Felis domesticus papillomavirus type 1

Fenneropenaeus chinensis hepatopancreatic densovirus

Figwort mosaic virus

Finch circovirus Finch polyomavirus

Flavobacterium phage 11b

Fowl adenovirus A

Fowl adenovirus D

Fowlpox virus

Francolinus leucoscepus papillomavirus 1

Fringilla coelebs papillomavirus

Frog adenovirus 1

Frog virus 3

Galleria mellonella densovirus

Gallid herpesvirus 1

Gallid herpesvirus 2

Gallid herpesvirus 3

Gammapapillomavirus HPV127

Geobacillus phage

GBSV1 Geobacillus virus E2

Glossina pallidipes salivary gland hypertrophy virus

Glypta fumiferanae ichnovirus segmentA10

Glypta fumiferanae ichnovirus segment A1

Glypta fumiferanae ichnovirus segment A2

Glypta fumiferanae ichnovirus segment A3

Glypta fumiferanae ichnovirus segment A4

Glypta fumiferanae ichnovirus segment A5

Glypta fumiferanae ichnovirus segment A6

Glypta fumiferanae ichnovirus segment A7

Glypta fumiferanae ichnovirus segment A8

Glypta fumiferanae ichnovirus segment A9

Glypta fumiferanae ichnovirus segment B10

Glypta fumiferanae ichnovirus segment B11

Glypta fumiferanae ichnovirus segment B12

Glypta fumiferanae ichnovirus segment B13

Glypta fumiferanae ichnovirus segment B14

Glypta fumiferanae ichnovirus segment B15

Glypta fumiferanae ichnovirus segment B16

Glypta fumiferanae ichnovirus segment B17

Glypta fumiferanae ichnovirus segment B18

Glypta fumiferanae ichnovirus segment B19

Glypta fumiferanae ichnovirus segment-B1

Glypta fumiferanae ichnovirus segment B20

Glypta fumiferanae ichnovirus segment B21

Glypta fumiferanae ichnovirus segment B22

Glypta fumiferanae ichnovirus segment B23

Glypta fumiferanae ichnovirus segment B24

Glypta fumiferanae ichnovirus segment B25

Glypta fumiferanae ichnovirus segment B26

Glypta fumiferanae ichnovirus segment B27

Glypta fumiferanae ichnovirus segment B28

Glypta fumiferanae ichnovirus segment B29

Glypta fumiferanae ichnovirus segment-B2

Glypta fumiferanae ichnovirus segment B30

Glypta fumiferanae ichnovirus segment B31

Glypta fumiferanae ichnovirus segment B32

Glypta fumiferanae ichnovirus segment B33

Glypta fumiferanae ichnovirus segment B34

Glypta fumiferanae ichnovirus segment B35

Glypta fumiferanae ichnovirus segment B36

Glypta fumiferanae ichnovirus segment B37

Glypta fumiferanae ichnovirus segment B38

Glypta fumiferanae ichnovirus segment B39

Glypta fumiferanae ichnovirus segment-B3

Glypta fumiferanae ichnovirus segment B40

Glypta fumiferanae ichnovirus segment B41

Glypta fumiferanae ichnovirus segment B42

Glypta fumiferanae ichnovirus segment B43

Glypta fumiferanae ichnovirus segment B44

Glypta fumiferanae ichnovirus segment B45

Glypta fumiferanae ichnovirus segment B46

Glypta fumiferanae ichnovirus segment B47

Glypta fumiferanae ichnovirus segment B48

Glypta fumiferanae ichnovirus segment B49

Glypta fumiferanae ichnovirus segment-B4

Glypta fumiferanae ichnovirus segment B50

Glypta fumiferanae ichnovirus segment B51

Glypta fumiferanae ichnovirus segment B52

Glypta fumiferanae ichnovirus segment B53

Glypta fumiferanae ichnovirus segment B54

Glypta fumiferanae ichnovirus segment B55

Glypta fumiferanae ichnovirus segment B56

Glypta fumiferanae ichnovirus segment B57

Glypta fumiferanae ichnovirus segment B58

Glypta fumiferanae ichnovirus segment B59

Glypta fumiferanae ichnovirus segment-B5

Glypta fumiferanae ichnovirus segment B60

Glypta fumiferanae ichnovirus segment B61

Glypta fumiferanae ichnovirus segment B62

Glypta fumiferanae ichnovirus segment B63

Glypta fumiferanae ichnovirus segment B64

Glypta fumiferanae ichnovirus segment B65

Glypta fumiferanae ichnovirus segment-B6

Glypta fumiferanae ichnovirus segment B7

Glypta fumiferanae ichnovirus segment B8

Glypta fumiferanae ichnovirus segment B9

Glypta fumiferanae ichnovirus segment C10

Glypta fumiferanae ichnovirus segment C11

Glypta fumiferanae ichnovirus segment C12

Glypta fumiferanae ichnovirus segment C13

Glypta fumiferanae ichnovirus segment C14

Glypta fumiferanae ichnovirus segment C15

Glypta fumiferanae ichnovirus segment C16

Glypta fumiferanae ichnovirus segment C17

Glypta fumiferanae ichnovirus segment C18

Glypta fumiferanae ichnovirus segment C19

Glypta fumiferanae ichnovirus segment-C1

Glypta fumiferanae ichnovirus segment C20

Glypta fumiferanae ichnovirus segment C21

Glypta fumiferanae ichnovirus segment C22

Glypta fumiferanae ichnovirus segment-C2

Glypta fumiferanae ichnovirus segment C3

Glypta fumiferanae ichnovirus segment C4

Glypta fumiferanae ichnovirus segment C5

Glypta fumiferanae ichnovirus segment C6

Glypta fumiferanae ichnovirus segment C7

Glypta fumiferanae ichnovirus segment C8

Glypta fumiferanae ichnovirus segment C9

Glypta fumiferanae ichnovirus segment D1

Glypta fumiferanae ichnovirus segment D2

Glypta fumiferanae ichnovirus segment D3

Glypta fumiferanae ichnovirus segment D4

Glypta fumiferanae ichnovirus segment D5

Glypta fumiferanae ichnovirus segment D6

Glypta fumiferanae ichnovirus segment D7

Glypta fumiferanae ichnovirus segment E1

Goatpox virus Pellor Goose circovirus

Goose hemorrhagic polyomavirus

Goose parvovirus

Gossypium darwinii symptomless alphasatellite DNA-alpha

Gossypium darwinii symptomless virus DNA-A

Gossypium davidsonii symptomless alphasatellite DNA-alpha-B

Gossypium mustilinum symptomless alphasatellite DNA-alpha-B

Gossypium punctatum mild leaf curl virus DNA A

Gossypium punctatum mild leaf curl virus DNA B

Ground squirrel hepatitis virus

Gryllus bimaculatus nudivirus Gull circovirus

Haemophilus phage HP1

Haemophilus phage HP2

Haloarcula hispanica pleomorphic virus 1

Haloarcula phage SH1

Halomonas phage phiHAP-1

Halorubrum phage HF2

Halorubrum pleomorphic virus 1

Halovirus HF1

Hamster polyomavirus

Helicoverpa armigera granulovirus

Helicoverpa armigera multiple nucleopolyhedrovirus

Helicoverpa armigera-NPV

Helicoverpa armigera NPV NNg1

Helicoverpa armigera nucleopolyhedrovirus G4

Helicoverpa zea SNPV

Heliothis virescens ascovirus 3e

Hepatitis B virus

Heron hepatitis B virus

His1 virus

His2 virus

Hollyhock leaf crumple virus

Honeysuckle yellow vein beta-JapanFukui2001

Honeysuckle yellow vein mosaic beta-JapanMiyizaki2001

Honeysuckle yellow vein mosaic disease associated satellite DNA beta-Ibaraki

Honeysuckle yellow vein mosaic Virus

Honeysuckle yellow vein mosaic virus-Kagoshima

Honeysuckle yellow vein mosaic virus satellite DNA beta

Honeysuckle yellow vein virus-UK1

Horsegram yellow mosaic Virus DNA B

Horsegram yellow mosaic virus

Horseradish curly top virus

Human adenovirus 54

Human adenovirus A

Human adenovirus B1

Human adenovirus B2

Human adenovirus C

Human adenovirus D

Human adenovirus E

Human adenovirus F

Human bocavirus 1

Human bocavirus 2

Human bocavirus 3

Human bocavirus 4

Human erythrovirus V9

Human herpesvirus 1

Human herpesvirus 2

Human herpesvirus 3

Human herpesvirus 4

Human herpesvirus 5 strain Merlin

Human herpesvirus 7

Human herpesvirus 8

Human papillomavirus-18

Human papillomavirus 1

Human papillomavirus-2

Human papillomavirus 54

Human papillomavirus-5

Human papillomavirus type 101

Human papillomavirus type 103

Human papillomavirus type 108

Human papillomavirus type-10

Human papillomavirus type 16

Human papillomavirus type 26

Human papillomavirus type 32

Human papillomavirus type 34

Human papillomavirus type 41

Human papillomavirus type 48

Human papillomavirus type 49

Human papillomavirus type-4

Human papillomavirus type 50

Human papillomavirus type 53

Human papillomavirus type 60

Human papillomavirus type 63

Human papillomavirus type 6b

Human papillomavirus type 7

Human papillomavirus type 88

Human papillomavirus type 90

Human papillomavirus type 92

Human papillomavirus type 96

Human papillomavirus type-9

Human parvovirus B19

Human T-lymphotropic virus 1

Human T-lymphotropic virus 4

Hyperthermophilic Archaeal Virus 1

Hyperthermophilic Archaeal Virus 2

Hyphantria cunea nucleopolyhedrovirus

Hyposoter fugitivus ichnovirus segment A1

Hyposoter fugitivus ichnovirus segment A2

Hyposoter fugitivus ichnovirus segment A3

Hyposoter fugitivus ichnovirus segment B10

Hyposoter fugitivus ichnovirus segment B11

Hyposoter fugitivus ichnovirus segment B12

Hyposoter fugitivus ichnovirus segment B13

Hyposoter fugitivus ichnovirus segment B14

Hyposoter fugitivus ichnovirus segment B15

Hyposoter fugitivus ichnovirus segment B16

Hyposoter fugitivus ichnovirus segment B17

Hyposoter fugitivus ichnovirus segment B18

Hyposoter fugitivus ichnovirus segment-B1

Hyposoter fugitivus ichnovirus segment B2

Hyposoter fugitivus ichnovirus segment B3

Hyposoter fugitivus ichnovirus segment B4

Hyposoter fugitivus ichnovirus segment B5

Hyposoter fugitivus ichnovirus segment B6

Hyposoter fugitivus ichnovirus segment B7

Hyposoter fugitivus ichnovirus segment B8

Hyposoter fugitivus ichnovirus segment B9

Hyposoter fugitivus ichnovirus segment C10

Hyposoter fugitivus ichnovirus segment C11

Hyposoter fugitivus ichnovirus segment C12

Hyposoter fugitivus ichnovirus segment C13

Hyposoter fugitivus ichnovirus segment C14

Hyposoter fugitivus ichnovirus segment C15

Hyposoter fugitivus ichnovirus segment C16

Hyposoter fugitivus ichnovirus segment C17

Hyposoter fugitivus ichnovirus segment C18

Hyposoter fugitivus ichnovirus segment C19

Hyposoter fugitivus ichnovirus segment-C1

Hyposoter fugitivus ichnovirus segment C20

Hyposoter fugitivus ichnovirus segment-C2

Hyposoter fugitivus ichnovirus segment C3

Hyposoter fugitivus ichnovirus segment C4

Hyposoter fugitivus ichnovirus segment C5

Hyposoter fugitivus ichnovirus segment C6

Hyposoter fugitivus ichnovirus segment C7

Hyposoter fugitivus ichnovirus segment C8

Hyposoter fugitivus ichnovirus segment C9

Hyposoter fugitivus ichnovirus segment D10

Hyposoter fugitivus ichnovirus segment D11

Hyposoter fugitivus ichnovirus segment D12

Hyposoter fugitivus ichnovirus segment-D1

Hyposoter fugitivus ichnovirus segment D2

Hyposoter fugitivus ichnovirus segment D3

Hyposoter fugitivus ichnovirus segment D4

Hyposoter fugitivus ichnovirus segment D5

Hyposoter fugitivus ichnovirus segment D6

Hyposoter fugitivus ichnovirus segment D7

Hyposoter fugitivus ichnovirus segment D8

Hyposoter fugitivus ichnovirus segment D9

Hyposoter fugitivus ichnovirus segment E1

Hyposoter fugitivus ichnovirus segment E2

Hyposoter fugitivus ichnovirus segment G1

Ictalurid herpesvirus 1 strain Auburn 1

Indian cassava mosaic virus DNA A

Indian cassava mosaic virus DNA B

Infectious hypodermal and hematopoietic necrosis virus

Infectious spleen and kidney necrosis virus

Invertebrate iridescent virus 6

Iodobacteriophage phiPLPE

Ipomoea yellow vein virus

Jatropha leaf curl virus DNA A

Jatropha yellow mosaic India virus DNA-A

JC polyomavirus

Junonia coenia densovirus

Kalanchoe top-spotting virus

Kenaf leaf curl virus DNA A

KI polyomavirus Stockholm 60

Klebsiella phage KP15

Klebsiella phage KP32

Klebsiella phage KP34

Klebsiella phage phiKO2

Kluyvera phage Kvp1

Kudzu mosaic virus DNA-A

Kudzu mosaic virus DNA-B

Lactobacillus johnsonii prophage Lj771

Lactobacillus phage A2

Lactobacillus phage KC5a

Lactobacillus phage Lb338-1

Lactobacillus phage Lc-Nu

Lactobacillus phage LL-H

Lactobacillus phage LP65

Lactobacillus phage Lrm1

Lactobacillus phage Lv-1

Lactobacillus phage phiAT3

Lactobacillus phage phig1e

Lactobacillus phage phiJL-1

Lactobacillus prophage Lj928

Lactobacillus prophage Lj965

Lactobacillus prophage phiadh

Lactococcus phage 1706

Lactococcus phage 4268

Lactococcus phage 712

Lactococcus phage asccphi28

Lactococcus phage bIBB29

Lactococcus phage bIL170

Lactococcus phage bIL67

Lactococcus phage BK5-T

Lactococcus phage c2

Lactococcus phage jj50

Lactococcus phage KSY1

Lactococcus phage P008

Lactococcus phage P087

Lactococcus phage phiLC3

Lactococcus phage Q54

Lactococcus phage r1t

Lactococcus phage sk1

Lactococcus phage TP901-1

Lactococcus phage Tuc2009

Lactococcus phage ul36

Lactococcus prophage bIL285

Lactococcus prophage bIL286

Lactococcus prophage bIL309

Lactococcus prophage bIL310

Lactococcus prophage bIL311

Lactococcus prophage bIL312

Lamium leaf distortion associated virus

Leucania separata nuclear polyhedrosis virus

Leucas zeylanica yellow vein virus satellite DNA beta

Lindernia anagallis yellow vein virus DNA-A

Lindernia anagallis yellow vein virus satellite DNA beta

Listeria phage A006

Listeria phage A118

Listeria phage A500

Listeria phage A511

Listeria phage B025

Listeria phage B054

Listeria phage P35

Listeria phage P40

Listonella phage phiHSIC

Loofa yellow mosaic virus DNA A

Loofa yellow mosaic virus DNA B

Lucky bamboo bacilliform virus

Ludwigia leaf distortion betasatellite IndiaAmadalavalasaHibiscus2 007

Ludwigia yellow vein virus-associated DNA beta

Ludwigia yellow vein virus DNA-A

Luffa begomovirus associated DNA beta

Luffa puckering and leaf distortion-associated DNA beta

LuIII virus

Lumpy skin disease virus NI-2490

Lymantria dispar MNPV

Lymantria xylina MNPV

Lymphocystis disease virus 1

Lymphocystis disease virus-isolate China

Macacine herpesvirus 1

Macacine herpesvirus 3

Macacine herpesvirus 4

Macacine herpesvirus 5

Macaque simian foamy virus

Macroptilium golden mosaic virus-

JamaicaWissadulaAugust Town DNA A

Macroptilium golden mosaic virus-

JamaicaWissadulaAugust Town DNA B

Macroptilium mosaic Puerto Rico virus DNA A

Macroptilium mosaic Puerto Rico virus DNA B

Macroptilium yellow mosaic Florida virus DNA A

Macroptilium yellow mosaic Florida virus DNA B

Macroptilium yellow mosaic virus DNA A

Macroptilium yellow mosaic virus DNA B

Maize streak virus-ASouth Africa

Malachra yellow vein mosaic virus-associated satellite DNA beta

Mal de Rio Cuarto virus segment 9

Malvastrum leaf curl Guangdong virus

Malvastrum leaf curl virus-associated defective DNA beta

Malvastrum leaf curl virus-G87

Malvastrum yellow mosaic virus-associated DNA 1

Malvastrum yellow mosaic virus DNA-A

Malvastrum yellow mosaic virus satellite DNA beta

Malvastrum yellow vein Baoshan virus DNA-A

Malvastrum yellow vein-virus

Malvastrum yellow vein virus satellite DNA beta

Malvastrum yellow vein Yunnan-virus

Malvastrum yellow vein Yunnan virus satellite DNA beta

Mamestra configurata NPV-A

Mamestra configurata NPV-B

Mannheimia phage phiMHaA1

Maruca vitrata MNPV

Mastomys coucha papillomavirus 2

Mastomys natalensis papillomavirus

Melanoplus sanguinipes entomopoxvirus

Meleagrid herpesvirus 1

Melon chlorotic leaf curl virus DNA A

Melon chlorotic mosaic virus-associated alphasatellite

Melon chlorotic mosaic virus DNA-A

Melon chlorotic mosaic virus DNA-B

Merkel cell polyomavirus

Merremia mosaic virus DNA A

Merremia mosaic virus DNA B

Mesta yellow vein mosaic Bahraich virus-

IndiaBahraich2007 DNA A

Mesta yellow vein mosaic virus-associated DNA beta

Mesta yellow vein mosaic virus DNA-A

Methanobacterium phage psiM2

Methanothermobacter prophage psiM100

Microbacterium phage Min1

Microcystis phage Ma-LMM01

Microplitis demolitor bracovirus segment A

Microplitis demolitor bracovirus segment B

Microplitis demolitor bracovirus segment C

Microplitis demolitor bracovirus segment D

Microplitis demolitor bracovirus segment E

Microplitis demolitor bracovirus segment F

Microplitis demolitor bracovirus segment G

Microplitis demolitor bracovirus segment H

Microplitis demolitor bracovirus segment I

Microplitis demolitor bracovirus segment J

Microplitis demolitor bracovirus segment K

Microplitis demolitor bracovirus segment L

Microplitis demolitor bracovirus segment M

Microplitis demolitor bracovirus segment N

Microplitis demolitor bracovirus segment O

Milk vetch dwarf virus segment 10

Milk vetch dwarf virus segment 11

Milk vetch dwarf virus segment-1

Milk vetch dwarf virus segment 2

Milk vetch dwarf virus segment 3

Milk vetch dwarf virus segment 4

Milk vetch dwarf virus segment 5

Milk vetch dwarf virus segment 6

Milk vetch dwarf virus segment 7

Milk vetch dwarf virus segment 8

Milk vetch dwarf virus segment 9

Mimosa yellow leaf curl virus-associated DNA 1

Mimosa yellow leaf curl virus DNA-A

Mimosa yellow leaf curl virus satellite DNA beta

Minute virus of mice

Mirabilis mosaic virus

Miscanthus streak virus-91

Molluscum contagiosum virus subtype 1

Monkeypox virus Zaire-96-I-16

Morganella phage MmP1

Mouse mammary tumor virus

Mouse parvovirus 1

Mouse parvovirus 2

Mouse parvovirus 3

Mouse parvovirus 4

Mouse parvovirus 5

Mulard duck circovirus

Mungbean yellow mosaic India virus DNA A

Mungbean yellow mosaic India virus DNA B

Mungbean yellow mosaic virus DNA A

Mungbean yellow mosaic virus DNA B

Murid herpesvirus 1

Murid herpesvirus 2

Murid herpesvirus 4

Murine adenovirus 3

Murine adenovirus A

Murine pneumotropic virus

Murine polyomavirus

Murine type C retrovirus

Musca domestica salivary gland hypertrophy virus

Muscovy duck circovirus

Muscovy duck parvovirus

Mus musculus papillomavirus type 1

Mycobacterium phage 244

Mycobacterium phage Adjutor

Mycobacterium phage Angel

Mycobacterium phage angelica

Mycobacterium phage Ardmore

Mycobacterium phage Barnyard

Mycobacterium phage Bethlehem

Mycobacterium phage Boomer

Mycobacterium phage BPs

Mycobacterium phage Brujita

Mycobacterium phage Butterscotch

Mycobacterium phage Bxb1

Mycobacterium phage Bxz1

Mycobacterium phage Bxz2

Mycobacterium phage Cali

Mycobacterium phage Catera

Mycobacterium phage Chah

Mycobacterium phage Che12

Mycobacterium phage Che8

Mycobacterium phage Che9c

Mycobacterium phage Che9d

Mycobacterium phage Cjw1

Mycobacterium phage Cooper

Mycobacterium phage Corndog

Mycobacterium phage CrimD

Mycobacterium phage D29

Mycobacterium phage DD5

Mycobacterium phage ET08

Mycobacterium phage Fruitloop

Mycobacterium phage Giles

Mycobacterium phage Gumball

Mycobacterium phage Halo

Mycobacterium phage Jasper

Mycobacterium phage KBG

Mycobacterium phage Konstantine

Mycobacterium phage Kostya

Mycobacterium phage L5

Mycobacterium phage LeBron

Mycobacterium phage Llij

Mycobacterium phage Lockley

Mycobacterium phage Myrna

Mycobacterium phage Nigel

Mycobacterium phage Omega

Mycobacterium phage Orion

Mycobacterium phage Pacc40

Mycobacterium phage PBI1

Mycobacterium phage Peaches

Mycobacterium phage PG1

Mycobacterium phage Phaedrus

Mycobacterium phage Phlyer

Mycobacterium phage Pipefish

Mycobacterium phage Plot

Mycobacterium phage PMC

Mycobacterium phage Porky

Mycobacterium phage Predator

Mycobacterium phage Pukovnik

Mycobacterium phage Qyrzula

Mycobacterium phage Ramsey

Mycobacterium phage Rizal

Mycobacterium phage Rosebush

Mycobacterium phage ScottMcG

Mycobacterium phage Solon

Mycobacterium phage Spud

Mycobacterium phage TM4

Mycobacterium phage Troll4

Mycobacterium phage Tweety

Mycobacterium phage U2

Mycobacterium phage Wildcat

Mycoplasma phage MAV1

Mycoplasma phage P1

Mycoplasma phage phiMFV1

Myotis polyomavirus VM-2008

Mythimna loreyi densovirus

Myxococcus phage Mx8

Myxoma virus

Myzus persicae densovirus

Nanovirus-like particle

Natrialba phage PhiCh1

Neodiprion abietis NPV

Neodiprion lecontei NPV

Neodiprion sertifer NPV Oat dwarf virus

OkLCV satDNA 10

Okra leaf curl disease associated DNA 1

Okra leaf curl Mali virus satellite DNA beta

Okra leaf curl virus-Cameroon

Okra mottle virus-Brazilokra DNA A

Okra mottle virus-Brazilokra DNA B

Okra yellow crinkle virus segment A

Okra yellow mosaic Mexico virus DNA A

Okra yellow mosaic Mexico virus DNA B

Okra yellow vein disease associated sequence virion

Okra yellow vein mosaic virus

Old World harvest mouse papillomavirus

Orangutan polyomavirus Orf virus

Orgyia leucostigma NPV

Orgyia pseudotsugata MNPV

Oryctes rhinoceros virus

Ostreid herpesvirus 1

Ostreococcus tauri virus 1

Ostreococcus virus OsV5

Ovine adenovirus A

Ovine adenovirus D

Ovine herpesvirus 2

Ovine papillomavirus-1

Panicum streak virus-Karino

Panine herpesvirus 2 strain Heberling

Papaya leaf curl China virus-G8

Papaya leaf curl China virus satellite DNA beta

Papaya leaf curl Guandong virus-GD2 DNA A

Papaya leaf curl virus-associated DNA beta

Papaya leaf curl-virus Papiine herpesvirus 2

Paramecium bursaria Chlorella virus 1

Paramecium bursaria Chlorella virus AR158

Paramecium bursaria Chlorella virus FR483

Paramecium bursaria Chlorella virus NY2A

Parvovirus H1

Passionfruit severe leaf distortion virus DNA-A

Passionfruit severe leaf distortion virus DNA-B

Pasteurella phage F108

Peanut chlorotic streak virus

Pedilanthus leaf curl virus-

Pedilanthus PakistanMultan2004

Pelargonium vein banding virus

Penaeus merguiensis densovirus

Penaeus monodon hepatopancreatic parvovirus

Pepper curly top virus

Pepper golden mosaic virus DNA A

Pepper golden mosaic virus DNA B

Pepper huasteco yellow vein virus DNA A

Pepper huasteco yellow vein virus DNA B

Pepper leaf curl Bangladesh virus segment A component

Pepper leaf curl virus DNA-A

Pepper leaf curl virus satellite DNA beta

Pepper leaf curl Yunnan virus satellite DNA beta

Pepper leaf curl Yunnan virus-YN323

Pepper yellow dwarf virus-New Mexico

Pepper yellow leaf curl Indonesia virus DNA-A

Pepper yellow leaf curl Indonesia virus DNA-B

Pepper yellow vein Mali virus

Periplaneta fuliginosa densovirus

Petunia vein clearing virus

Phage cdtI

Phage Gifsy-1

Phage Gifsy-2

Phage phiJL001

Phocoena spinipinnis papillomavirus

Phormidium phage Pf-WMP3

Phormidium phage Pf-WMP4

Phthorimaea operculella granulovirus

Pieris rapae granulovirus

Planaria asexual strain-specific virus-like element type 1 large DNA segment

Planaria asexual strain-specific virus-like element type 1 small DNA segment

Planococcus citri densovirus

Plutella xylostella granulovirus

Plutella xylostella multiple nucleopolyhedrovirus

Polyomavirus HPyV6

Polyomavirus HPyV7

Porcine adenovirus C

Porcine circovirus 1

Porcine circovirus 2

Porcine endogenous retrovirus E

Porcine parvovirus

Potato apical leaf curl disease-associated satellite DNA beta

Potato yellow mosaic

Panama virus DNA A

Potato yellow mosaic

Panama virus DNA B

Potato yellow mosaic Trinidad virus DNA A

Potato yellow mosaic Trinidad virus DNA B

Potato yellow mosaic virus DNA A

Potato yellow mosaic virus DNA B

Prochlorococcus phage P-SSM4

Propionibacterium phage B5

Propionibacterium phage PA6

Pseudaletia unipuncta granulovirus

Pseudoalteromonas phage PM2

Pseudocowpox virus

Pseudomonas phage 119X

Pseudomonas phage 14-1

Pseudomonas phage 201phi2-1

Pseudomonas phage 73

Pseudomonas phage B3

Pseudomonas phage D3112

Pseudomonas phage-D3

Pseudomonas phage DMS3

Pseudomonas phage EL

Pseudomonas phage F10

Pseudomonas phage F116

Pseudomonas phage F8

Pseudomonas phage gh-1

Pseudomonas phage LBL3

Pseudomonas phage LIT1

Pseudomonas phage LKD16

Pseudomonas phage LMA2

Pseudomonas phage LUZ19

Pseudomonas phage LUZ24

Pseudomonas phage LUZ7

Pseudomonas phage M6

Pseudomonas phage MP22

Pseudomonas phage MP29

Pseudomonas phage MP38

Pseudomonas phage PA11

Pseudomonas phage PAJU2

Pseudomonas phage PaP2

Pseudomonas phage PaP3

Pseudomonas phage PB1

Pseudomonas phage Pf1

Pseudomonas phage Pf3

Pseudomonas phage phi-2

Pseudomonas phage phiCTX

Pseudomonas phage phikF77

Pseudomonas phage phiKMV

Pseudomonas phage phiKZ

Pseudomonas phage PT2

Pseudomonas phage PT5

Pseudomonas phage SN

Pseudomonas phage YuA

Psittacid herpesvirus 1

Psittacus erithacus timneh papillomavirus

Pumpkin yellow mosaic Malaysia virus DNA A

Pyrobaculum spherical virus

Pyrococcus abyssi virus 1

Rabbit fibroma virus

Rabbit oral papillomavirus

Rachiplusia ou MNPV

Radish leaf curl virus satellite DNA beta

Radish leaf curl virus segment A

Ralstonia phage p12J

Ralstonia phage phiRSA1

Ralstonia phage RSB1

Ralstonia phage RSL1

Ralstonia phage RSM1

Ralstonia phage RSM3

Ralstonia phage RSS1

Ramie mosaic virus DNA-A

Ramie mosaic virus DNA-B

Ranid herpesvirus 1 strain McKinnell

Ranid herpesvirus 2 strain ATCC VR-568

Rauscher murine leukemia virus

Raven circovirus RD114 retrovirus

Reticuloendotheliosis virus

Rhesus monkey

Papillomavirus

Rhizobium phage 16-3

Rhodothermus phage RM378

Rhynchosia golden mosaic virus DNA A

Rhynchosia golden mosaic virus DNA B

Rhynchosia golden mosaic Yucatan virus DNA A

Rhynchosia golden mosaic Yucatan virus DNA B

Rice tungro bacilliform virus

Roseobacter phage SIO1

Roseophage DSS3P2

Roseophage EE36P1

Rosss goose hepatitis B virus

Rousettus aegyptiacus papillomavirus type 1

Rudbeckia flower distortion virus

Saccharum streak virus

Saimiriine herpesvirus 2

Salmonella enterica bacteriophage SE1

Salmonella phage c341

Salmonella phage E1

Salmonella phage epsilon34

Salmonella phage Fels-1

Salmonella phage phiSG-JL2

Salmonella phage SETP3

Salmonella phage ST64B

Sclerotinia sclerotiorum hypovirulence associated DNA virus 1

Sea turtle tornovirus 1

Senecio yellow mosaic virus

Sheeppox virus 17077-99

Sheldgoose hepatitis B virus

Shigella phage phiSboM-AG3

Shrimp white spot syndrome virus

Sida golden mosaic Costa Rica virus DNA A

Sida golden mosaic Costa Rica virus DNA B

Sida golden mosaic Florida virus-Malvastrum DNA-A

Sida golden mosaic Florida virus-Malvastrum DNA-B

Sida golden mosaic Honduras virus DNA A

Sida golden mosaic Honduras virus DNA B

Sida golden mosaic virus DNA-A

Sida golden mosaic virus DNA-B

Sida golden mottle virus DNA-A

Sida golden mottle virus DNA-B

Sida leaf curl virus-associated DNA 1

Sida leaf curl virus-associated DNA beta

Sida leaf curl-virus

Sida leaf curl virus satellite DNA beta

Sida micrantha mosaic virus segment A

Sida micrantha mosaic virus segment B

Sida mosaic Sinaloa virus DNA A

Sida mosaic Sinaloa virus DNA B

Sida mottle virus

Sida yellow mosaic virus-China-associated DNA beta DNA beta

Sida yellow mosaic-virus

Sida yellow mosaic Yucatan virus DNA A

Sida yellow mosaic Yucatan virus DNA B

Sida yellow vein disease associated DNA 1

Sida yellow vein Madurai virus

Sida yellow vein Vietnam virus-associated DNA 1

Sida yellow vein Vietnam virus DNA-A

Sida yellow vein Vietnam virus satellite DNA beta

Sida yellow vein virus DNA A

Sida yellow vein virus DNA B

Sida yellow vein virus satellite DNA beta

Siegesbeckia yellow vein Guangxi virus

Siegesbeckia yellow vein virus-GD13-associated DNA beta

Siegesbeckia yellow vein virus GD13

Simian adenovirus 1

Simian adenovirus 3

Simian foamy virus 3

Simian immunodeficiency virus SIV-mnd 2

Simian retrovirus 4

Simian T-cell lymphotropic virus 6

Simian T-lymphotropic virus 1

Simian T-lymphotropic virus 3

Simian virus 40

Singapore grouper iridovirus

Sinorhizobium phage PBC5

Small anellovirus 1

Small anellovirus 2

Snake parvovirus 1

Snow goose hepatitis B virus

Sodalis phage phiSG1

Sodalis phage SO-1

South African cassava mosaic virus DNA A

South African cassava mosaic virus DNA B

Soybean chlorotic blotch virus DNA A

Soybean chlorotic blotch virus DNA B

Soybean chlorotic mottle virus

Soybean crinkle leaf virus

Soybean mild mottle virus

Spilanthes yellow vein virus DNA-A

Spinach curly top virus

Spiroplasma kunkelii virus

SkV1 CR2-3x

Spiroplasma phage 1-C74

Spiroplasma phage 1-R8A2B

Spiroplasma phage 4

Spiroplasma phage SVTS2

Spodoptera exigua MNPV

Spodoptera frugiperda ascovirus 1a

Spodoptera frugiperda MNPV virus

Spodoptera litura granulovirus

Spodoptera litura NPV

Spodoptera litura nucleopolyhedrovirus II

Sputnik virophage

Squash leaf curl China virus-B DNA-A

Squash leaf curl China virus-B DNA B

Squash leaf curl Philippines virus segment A

Squash leaf curl Philippines virus segment B

Squash leaf curl virus A component DNA

Squash leaf curl virus B component DNA

Squash leaf curl Yunnan virus

Squash mild leaf curl virus-Imperial Valley DNA A

Squash mild leaf curl virus-Imperial Valley DNA B

Squash yellow mild mottle virus DNA B

Squirrel monkey polyomavirus

Sri Lankan cassava mosaic virus DNA A

Sri Lankan cassava mosaic virus DNA B

Stachytarpheta leaf curl virus

Staphylococcus phage 11

Staphylococcus phage 187

Staphylococcus phage 2638A

Staphylococcus phage 29

Staphylococcus phage 37

Staphylococcus phage 3A

Staphylococcus phage 42e

Staphylococcus phage 44AHJD

Staphylococcus phage 47

Staphylococcus phage 52A

Staphylococcus phage 53

Staphylococcus phage 55

Staphylococcus phage 66

Staphylococcus phage 69

Staphylococcus phage 71

Staphylococcus phage 77

Staphylococcus phage 80alpha

Staphylococcus phage 85

Staphylococcus phage 88

Staphylococcus phage 92

Staphylococcus phage 96

Staphylococcus phage CNPH82

Staphylococcus phage EW

Staphylococcus phage G1

Staphylococcus phage K

Staphylococcus phage P954

Staphylococcus phage PH15

Staphylococcus phage phi2958PVL

Staphylococcus phage phiETA2

Staphylococcus phage phiETA3

Staphylococcus phage-phiETA

Staphylococcus phage phiMR11

Staphylococcus phage phiMR25

Staphylococcus phage phiNM1

Staphylococcus phage phiNM3

Staphylococcus phage phiP68

Staphylococcus phage phiPVL108

Staphylococcus phage phiPVL-CN125

Staphylococcus phage phiSauS-IPLA35

Staphylococcus phage phiSauS-IPLA88

Staphylococcus phage phiSLT

Staphylococcus phage PT1028

Staphylococcus phage ROSA

Staphylococcus phage SAP-26

Staphylococcus phage SAP2

Staphylococcus phage Twort

Staphylococcus phage X2

Staphylococcus prophage phi 12

Staphylococcus prophage phi 13

Staphylococcus prophage phiN315

Staphylococcus prophage phiPV83

Staphylococcus prophage PVL

Staphylococcus prophage tp310-1

Staphylococcus prophage tp310-2

Staphylococcus prophage tp310-3

Starling circovirus

Stenotrophomonas phage phiSMA9

Stenotrophomonas phage S1

Strawberry vein banding virus

Streptococcus phage 2972

Streptococcus phage 5093

Streptococcus phage 7201

Streptococcus phage 858

Streptococcus phage Abc2

Streptococcus phage ALQ13.2

Streptococcus phage C1

Streptococcus phage Cp-1

Streptococcus phage DT1

Streptococcus phage M102

Streptococcus phage O1205

Streptococcus phage P9

Streptococcus phage PH10

Streptococcus phage PH15

Streptococcus phage phi3396

Streptococcus phage Sfi11

Streptococcus phage Sfi19

Streptococcus phage Sfi21

Streptococcus phage SM1

Streptococcus phage SMP

Streptococcus prophage 315.1

Streptococcus prophage 315.2

Streptococcus prophage 315.5

Streptococcus prophage 315.6

Streptococcus prophage EJ-1

Streptococcus prophage MM1

Streptococcus pyogenes phage 315.3

Streptomyces phage mu16

Streptomyces phage phiBT1

Streptomyces phage phiC31

Streptomyces phage phiSASD1

Streptomyces phage VWB Stx1 converting phage

Stx2-converting phage 1717

Stx2-converting phage 86

Stx2 converting phage-I

Stx2 converting phage II

Subterranean clover stunt virus DNA 1

Subterranean clover stunt virus DNA 2

Subterranean clover stunt virus DNA 3

Subterranean clover stunt virus DNA 4

Subterranean clover stunt virus DNA 5

Subterranean clover stunt virus DNA 6

Subterranean clover stunt virus DNA 7

Subterranean clover stunt virus DNA 8

Sugarcane bacilliform IM virus

Sugarcane bacilliform Mor virus

Sugarcane bacilliform virus

Sugarcane streak Egypt virus-Giza

Sugarcane streak Reunion virus

Sugarcane streak virus-Natal

Suid herpesvirus 1

Sulfolobus islandicus filamentous virus

Sulfolobus islandicus rod-shaped virus 1

Sulfolobus islandicus rod-shaped virus 2

Sulfolobus spindle-shaped virus 4

Sulfolobus spindle-shaped virus 5

Sulfolobus spindle-shaped virus 6

Sulfolobus spindle-shaped virus 7

Sulfolobus turreted icosahedral virus 2

Sulfolobus turreted icosahedral-virus

Sulfolobus virus 1

Sulfolobus virus 2

Sulfolobus virus Kamchatka 1

Sulfolobus virus Ragged Hills

Sulfolobus virus STSV1

Sunn hemp leaf distortion virus DNA-A

Sus scrofa papillomavirus type 1

Sweetpotato badnavirus B

Sweet potato leaf curl Bengal virus-IndiaWest Bengal2008 segment A

Sweet potato leaf curl Canary virus

Sweet potato leaf curl Georgia virus

Sweet potato leaf curl Lanzarote virus

Sweet potato leaf curl Spain virus

Sweet potato leaf curl virus

Swinepox virus Synechococcus phage P60

Synechococcus phage S-PM2

Synechococcus phage S-RSM4

Synechococcus phage syn9

Tanapox virus

Taro bacilliform virus

Taterapox virus

Thalassomonas phage BA3

Thermoproteus tenax spherical virus 1

Thermus phage IN93

Thermus phage P23-45

Thermus phage P23-77

Thermus phage P74-26

Thermus phage phiYS40

Tobacco curly shoot virus associated DNA 1

Tobacco curly shoot virus-associated DNA beta

Tobacco curly shoot-virus

Tobacco leaf curl disease associated sequence virion

Tobacco leaf curl Japan virus

Tobacco leaf curl Kochi virus

Tobacco leaf curl Thailand virus

Tobacco leaf curl virus-associated DNA beta

Tobacco leaf curl Yunnan virus associated DNA 1

Tobacco leaf curl Yunnan virus satellite DNA beta

Tobacco leaf curl Yunnan virus-Y136

Tobacco leaf curl Zimbabwe virus

Tobacco vein clearing virus

Tobacco yellow dwarf virus

Tomato begomovirus satellite DNA beta

Tomato chino La Paz virus segment A

Tomato chlorotic mottle virus DNA A

Tomato chlorotic mottle virus DNA B

Tomato common mosaic virus DNA-A

Tomato common mosaic virus DNA-B

Tomato curly stunt virus

Tomato golden mosaic virus DNA A

Tomato golden mosaic virus DNA B

Tomato golden mottle virus DNA A

Tomato golden mottle virus DNA B

Tomato leaf curl Arusha virus DNA-A

Tomato leaf curl Bangalore virus-Ban5 satellite DNA beta

Tomato leaf curl Bangalore-virus

Tomato leaf curl Bangladesh virus

Tomato leaf curl Cameroon virus-CameroonBueaOkra2008

Tomato leaf curl Cebu virus DNA-A

Tomato leaf curl China virus-G32

Tomato leaf curl Cotabato virus DNA-A

Tomato leaf curl Ghana virus segment A

Tomato leaf curl Guangdong virus DNA-A

Tomato leaf curl Guangxi virus

Tomato leaf curl Gujarat virus-Varanasi segment A

Tomato leaf curl Gujarat virus-Varanasi segment B

Tomato leaf curl Hainan virus

Tomato leaf curl Hsinchu virus-TaiwanHsinchu2005 DNA A

Tomato leaf curl Iran virus

Tomato leaf curl Java virus-Ageratum satellite DNA

Tomato leaf curl Java-virus

Tomato leaf curl Joydebpur beta virus

Tomato leaf curl Joydebpur virus DNA-A

Tomato leaf curl Karnataka virus-associated DNA beta DNA-A

Tomato leaf curl Karnataka-virus

Tomato leaf curl Kerala virus

Tomato leaf curl Kumasi virus segment A

Tomato leaf curl Laos virus

Tomato leaf curl Malaysia virus

Tomato leaf curl Mali virus

Tomato leaf curl Mayotte virus

Tomato leaf curl Mindanao virus DNA-A

Tomato leaf curl New Delhi virus-associated DNA beta

Tomato leaf curl New Delhi virus DNA A

Tomato leaf curl New Delhi virus DNA B

Tomato leaf curl Nigeria virus-Nigeria2006

Tomato leaf curl Pakistan virus associated DNA 1

Tomato leaf curl Pakistan virus segment A

Tomato leaf curl Palampur virus

Tomato leaf curl Patna virus DNA-A

Tomato leaf curl Philippines virus

Tomato leaf curl Philippine virus satellite DNA beta

Tomato leaf curl Pune virus

Tomato leaf curl Seychelles virus

Tomato leaf curl Sinaloa virus DNA A

Tomato leaf curl Sinaloa virus DNA B

Tomato leaf curl Sri Lanka virus

Tomato leaf curl Sudan virus-Gezira

Tomato leaf curl Sulawesi virus DNA-A

Tomato leaf curl Taiwan virus

Tomato leaf curl Togo virus-Togo2006

Tomato leaf curl Vietnam virus DNA A

Tomato leaf curl virus-associated DNA beta

Tomato leaf curl-virus

Tomato leaf curl virus-Pune-associated DNA beta DNA-A

Tomato mild mosaic virus DNA-A

Tomato mild mosaic virus DNA-B

Tomato mild yellow leaf curl Aragua virus DNA A

Tomato mild yellow leaf curl Aragua virus DNA B

Tomato mosaic Havana virus DNA A

Tomato mosaic Havana virus DNA B

Tomato mosaic leaf curl virus DNA A

Tomato mosaic leaf curl virus DNA B

Tomato mottle Taino virus DNA A

Tomato mottle Taino virus DNA B

Tomato mottle virus DNA A

Tomato mottle virus DNA B

Tomato pseudo-curly top virus

Tomato rugose mosaic virus DNA A

Tomato rugose mosaic virus DNA B

Tomato severe leaf curl virus

Tomato severe rugose virus DNA A

Tomato severe rugose virus DNA B

Tomato yellow dwarf disease associated satellite DNA beta-Kochi virus

Tomato yellow leaf curl China virus associated DNA 1

Tomato yellow leaf curl China-virus

Tomato yellow leaf curl Guangdong virus DNA-A

Tomato yellow leaf curl Indonesia virus-Lembang

Tomato yellow leaf curl Kanchanaburi virus DNA A

Tomato yellow leaf curl Kanchanaburi virus DNA B

Tomato yellow leaf curl Mali virus-associated DNA beta

Tomato yellow leaf curl Thailand virus associated DNA 1

Tomato yellow leaf curl Thailand virus DNA A

Tomato yellow leaf curl Thailand virus DNA B

Tomato yellow leaf curl Vietnam virus DNA-A

Tomato yellow leaf curl Vietnam virus satellite DNA beta

Tomato yellow leaf curl virus-associated DNA beta

Tomato yellow margin leaf curl virus DNA A

Tomato yellow margin leaf curl virus DNA B

Tomato yellow spot virus DNA-A

Tomato yellow spot virus DNA-B

Tomato yellow vein streak virus DNA-A

Tomato yellow vein streak virus DNA-B

Torque teno canis virus Torque teno douroucouli virus

Torque teno felis virus

Torque teno midi virus 1

Torque teno midi virus 2

Torque teno mini virus 1

Torque teno mini virus 2

Torque teno mini virus 3

Torque teno mini virus 4

Torque teno mini virus 5

Torque teno mini virus 6

Torque teno mini virus 7

Torque teno mini virus 8

Torque teno mini virus 9

Torque teno sus virus 1

Torque teno tamarin virus

Torque teno virus 10

Torque teno virus 12

Torque teno virus 14

Torque teno virus 15

Torque teno virus 16

Torque teno virus 19

Torque teno virus-1

Torque teno virus 25

Torque teno virus 26

Torque teno virus 27

Torque teno virus 28

Torque teno virus 3

Torque teno virus 4

Torque teno virus 6

Torque teno virus 7

Torque teno virus 8

Trichodysplasia spinulosa-associated polyomavirus

Trichoplusia ni ascovirus 2c

Trichoplusia ni SNPV

Tupaiid herpesvirus 1

Turkey adenovirus A

Turnip curly top virus

TYLCCNV-Y322 satellite DNA beta

Urochloa streak virus Vaccinia virus Variola virus

Velvet bean severe mosaic virus DNA A

Velvet bean severe mosaic virus DNA B

Vernonia yellow vein betasatellite

Vernonia yellow vein virus DNA-A

Vibrio phage fs1

Vibrio phage fs2

Vibrio phage K139

Vibrio phage kappa

Vibrio phage KSF-1phi

Vibrio phage KVP40

Vibrio phage N4

Vibrio phage VEJphi

Vibrio phage Vf12

Vibrio phage VfO3K6

Vibrio phage VfO4K68

Vibrio phage VGJphi

Vibrio phage VHML

Vibrio phage VP2

Vibriophage VP4

Vibrio phage VP5

Vibrio phage VP882

Vibrio phage VP93

Vibrio phage VpV262

Vibrio phage VSK

Watermelon chlorotic stunt virus DNA A

Watermelon chlorotic stunt virus DNA B

Wheat dwarf virus

Wissadula golden mosaic St Thomas Virus DNA A

Wissadula golden mosaic St Thomas Virus DNA B

Woodchuck hepatitis virus

WU Polyomavirus

Xanthomonas phage Cf1c

Xanthomonas phage OP1

Xanthomonas phage OP2

Xanthomonas phage phiL7

Xanthomonas phage Xop411

Xanthomonas phage Xp10

Xanthomonas phage Xp15

Xenopus laevis endogenous retrovirus Xen1

Xestia c-nigrum granulovirus

Xylella phage Xfas53

Yaba-like disease virus

Yaba monkey tumor virus

Yersinia pestis phage phiA1122

Yersinia phage Berlin

Yersinia phage L-413C

Yersinia phage phiYeO3-12

Yersinia phage PY54

Zinnia leaf curl disease associated sequence virion

Zinnia leaf curl virus-associated DNA beta

## Appendix 3: Bacterial Database

Acaryochloris marina MBIC11017

Acetobacter pasteurianus IFO 3283 01

Acetohalobium arabaticum DSM 5501

Acholeplasma laidlawii PG 8A

Achromobacter xylosoxidans A8

Acidaminococcus fermentans DSM 20731

Acidaminococcus intestini RyC MR95

Acidianus hospitalis W1

Acidilobus saccharovorans 345 15

Acidimicrobium ferrooxidans DSM 10331

Acidiphilium cryptum JF 5

Acidiphilium multivorum AIU301

Acidithiobacillus caldus SM 1

Acidithiobacillus ferrivorans SS3

Acidithiobacillus ferrooxidans ATCC 23270

Acidobacterium capsulatum ATCC 51196

Acidothermus cellulolyticus 11B

Acidovorax avenae subsp avenae ATCC 19860

Acidovorax citrulli AAC00 1

Acidovorax ebreus TPSY

Acidovorax sp JS42

Aciduliprofundum boonei T469

Acinetobacter baumannii 1656 2

Acinetobacter calcoaceticus PHEA 2

Acinetobacter oleivorans DR1

Acinetobacter sp ADP1

Actinobacillus pleuropneumoniae serovar 5b str L20

Actinobacillus succinogenes 130Z

Actinoplanes sp SE50 110

Actinosynnema mirum DSM 43827

Aerococcus urinae ACS 120 V Col10a

Aeromonas hydrophila subsp hydrophila ATCC 7966

Aeromonas salmonicida subsp salmonicida A449

Aeromonas veronii B565

Aeropyrum pernix K1

Aggregatibacter actinomycetemcomitans D11S 1

Aggregatibacter aphrophilus NJ8700

Agrobacterium fabrum str C58

Agrobacterium radiobacter K84

Agrobacterium sp H13 3

Agrobacterium vitis S4

Akkermansia muciniphila ATCC BAA 835

Alcanivorax borkumensis SK2

Alicycliphilus denitrificans BC

Alicyclobacillus acidocaldarius subsp acidocaldarius DSM 446

Aliivibrio salmonicida LFI1238

Alkalilimnicola ehrlichii MLHE 1

Alkaliphilus metalliredigens QYMF

Alkaliphilus oremlandii OhILAs

Allochromatium vinosum DSM 180

Alteromonas macleodii str Deep ecotype

Alteromonas sp SN2

Aminobacterium colombiense DSM 12261

Ammonifex degensii KC4

Amycolatopsis mediterranei U32

Amycolicicoccus subflavus DQS3 9A1

Anabaena variabilis ATCC 29413

Anaerococcus prevotii DSM 20548

Anaerolinea thermophila UNI 1

Anaeromyxobacter dehalogenans 2CP 1

Anaeromyxobacter sp Fw109 5

Anaplasma centrale str Israel

Anaplasma marginale str Florida

Anaplasma phagocytophilum HZ

Anoxybacillus flavithermus WK1

Aquifex aeolicus VF5

Arcanobacterium haemolyticum DSM 20595

Archaeoglobus fulgidus DSM 4304

Archaeoglobus profundus DSM 5631

Archaeoglobus veneficus SNP6

Arcobacter butzleri RM4018

Arcobacter nitrofigilis DSM 7299

Arcobacter sp L

Aromatoleum aromaticum EbN1

Arthrobacter arilaitensis Re117

Arthrobacter aurescens TC1

Arthrobacter chlorophenolicus A6

Arthrobacter phenanthrenivorans Sphe3

Arthrobacter sp FB24

Aster yellows witches broom phytoplasma AYWB

Asticcacaulis excentricus CB 48

Atopobium parvulum DSM 20469

Azoarcus sp BH72

Azorhizobium caulinodans ORS 571

Azospirillum sp B510

Azotobacter vinelandii DJ Bacillus amyloliquefaciens DSM 7

Bacillus anthracis str Ames

Bacillus atrophaeus 1942

Bacillus cellulosilyticus DSM 2522

Bacillus cereus 03BB102

Bacillus clausii KSM K16

Bacillus coagulans 2 6

Bacillus cytotoxicus NVH 391 98

Bacillus halodurans C 125

Bacillus licheniformis DSM 13 ATCC 14580

Bacillus megaterium DSM 319

Bacillus mycoides DSM 2048

Bacillus pseudofirmus OF4

Bacillus pumilus SAFR 032

Bacillus selenitireducens MLS10

Bacillus subtilis BSn5

Bacillus thuringiensis serovar berliner ATCC 10792

Bacillus weihenstephanensis KBAB4

Bacteriovorax marinus SJ

Bacteroides fragilis YCH46

Bacteroides helcogenes P 36 108

Bacteroides salanitronis DSM 18170

Bacteroides thetaiotaomicron VPI 5482

Bacteroides vulgatus ATCC 8482

Bartonella bacilliformis KC583

Bartonella clarridgeiae 73

Bartonella grahamii as4aup

Bartonella henselae str Houston 1

Bartonella quintana str Toulouse

Bartonella tribocorum CIP 105476

Baumannia cicadellinicola str Hc Homalodisca coagulata

Bdellovibrio bacteriovorus HD100

Beijerinckia indica subsp indica ATCC 9039

Beutenbergia cavernae DSM 12333

Bifidobacterium adolescentis ATCC 15703

Bifidobacterium animalis subsp animalis ATCC 25527

Bifidobacterium bifidum S17

Bifidobacterium breve ACS 071 V Sch8b

Bifidobacterium dentium Bd1

Bifidobacterium longum DJO10A

Blattabacterium sp

Blattella germanica str Bge

Bordetella avium 197N

Bordetella bronchiseptica RB50

Bordetella parapertussis 12822

Bordetella pertussis CS

Bordetella petrii DSM 12804

Borrelia afzelii PKo

Borrelia bissettii DN127

Borrelia burgdorferi B31

Borrelia duttonii Ly

Borrelia garinii PBi

Borrelia hermsii DAH

Borrelia recurrentis A1

Borrelia turicatae 91E135

Brachybacterium faecium DSM 4810

Brachyspira hyodysenteriae WA1

Brachyspira intermedia PWS A

Brachyspira murdochii DSM 12563

Brachyspira pilosicoli 95 1000

Bradyrhizobium japonicum USDA 110

Bradyrhizobium sp BTAi1

Brevibacillus brevis NBRC 100599

Brevundimonas subvibrioides ATCC 15264

Brucella abortus A13334

Brucella canis ATCC 23365

Brucella melitensis bv 1 str 16M

Brucella microti CCM 4915

Brucella ovis ATCC 25840

Brucella pinnipedialis B2 94

Brucella suis 1330

Buchnera aphidicola str APS Acyrthosiphon pisum

Burkholderia ambifaria AMMD

Burkholderia cenocepacia HI2424

Burkholderia gladioli BSR3

Burkholderia glumae BGR1

Burkholderia mallei ATCC 23344

Burkholderia multivorans ATCC 17616

Burkholderia phymatum STM815

Burkholderia phytofirmans PsJN

Burkholderia pseudomallei 668

Burkholderia rhizoxinica HKI 454

Burkholderia sp 383

Burkholderia thailandensis E264

Burkholderia vietnamiensis G4

Burkholderia xenovorans LB400

Butyrivibrio proteoclasticus B316

Caldicellulosiruptor bescii DSM 6725

Caldicellulosiruptor hydrothermalis 108

Caldicellulosiruptor kristjanssonii 177R1B

Caldicellulosiruptor kronotskyensis 2002

Caldicellulosiruptor lactoaceticus 6A

Caldicellulosiruptor obsidiansis OB47

Caldicellulosiruptor owensensis OL

Caldicellulosiruptor saccharolyticus DSM 8903

Calditerrivibrio nitroreducens DSM 19672

Caldivirga maquilingensis IC 167

Campylobacter concisus 13826

Campylobacter curvus 52592

Campylobacter fetus subsp fetus 82 40

Campylobacter hominis ATCC BAA 381

Campylobacter jejuni subsp jejuni 81116

Campylobacter lari RM2100

Candidatus Accumulibacter phosphatis clade IIA str UW 1

Candidatus Amoebophilus asiaticus 5a2

Candidatus Arthromitus sp SFB mouse Japan

Candidatus Azobacteroides pseudotrichonymphae genomovar CFP2

Candidatus Blochmannia floridanus

Candidatus Carsonella ruddii PV

Candidatus Chloracidobacterium thermophilum B

Candidatus Desulforudis audaxviator MP104C

Candidatus Hamiltonella defensa 5AT Acyrthosiphon pisum

Candidatus Hodgkinia cicadicola Dsem

Candidatus Korarchaeum cryptofilum OPF8

Candidatus Koribacter versatilis Ellin345

Candidatus Liberibacter asiaticus str psy62

Candidatus Methylomirabilis oxyfera

Candidatus Midichloria mitochondrii IricVA

Candidatus Moranella endobia PCIT

Candidatus Nitrospira defluvii

Candidatus Pelagibacter sp IMCC9063

Candidatus Phytoplasma australiense

Candidatus Protochlamydia amoebophila UWE25

Candidatus Puniceispirillum marinum IMCC1322

Candidatus Riesia pediculicola USDA

Candidatus Ruthia magnifica str Cm

Calyptogena magnifica

Candidatus Solibacter usitatus Ellin6076

Candidatus Sulcia muelleri GWSS

Candidatus Tremblaya princeps PCIT

Candidatus Vesicomyosocius okutanii HA

Candidatus Zinderia insecticola CARI

Capnocytophaga canimorsus Cc5

Capnocytophaga ochracea DSM 7271

Carboxydothermus hydrogenoformans Z 2901

Carnobacterium sp 17 4

Catenulispora acidiphila DSM 44928

Caulobacter crescentus CB15

Caulobacter segnis ATCC 21756

Caulobacter sp K31

Cellulomonas fimi ATCC 484

Cellulomonas flavigena DSM 20109

Cellulophaga algicola DSM 14237

Cellulophaga lytica DSM 7489

Cellvibrio gilvus ATCC 13127

Cellvibrio japonicus Ueda107

Cenarchaeum symbiosum A

Chelativorans sp BNC1

Chitinophaga pinensis DSM 2588

Chlamydia muridarum Nigg

Chlamydia trachomatis 434 Bu

Chlamydophila abortus S26 3

Chlamydophila caviae GPIC

Chlamydophila felis Fe C 56

Chlamydophila pecorum E58

Chlamydophila pneumoniae CWL029

Chlamydophila psittaci 6BC

Chlorobaculum parvum NCIB 8327

Chlorobium chlorochromatii CaD3

Chlorobium limicola DSM 245

Chlorobium luteolum DSM 273

Chlorobium phaeobacteroides DSM 266

Chlorobium phaeovibrioides DSM 265

Chlorobium tepidum TLS

Chloroflexus aggregans DSM 9485

Chloroflexus aurantiacus J 10 fl

Chloroflexus sp Y 400 fl

Chloroherpeton thalassium ATCC 35110

Chromobacterium violaceum ATCC 12472

Chromohalobacter salexigens DSM 3043

Citrobacter koseri ATCC BAA 895

Citrobacter rodentium ICC168

Clavibacter michiganensis subsp michiganensis NCPPB 382

Clostridium acetobutylicum ATCC 824

Clostridium beijerinckii NCIMB 8052

Clostridium botulinum A str ATCC 3502

Clostridium cellulolyticum H10

Clostridium cellulovorans 743B

Clostridium difficile 630

Clostridium kluyveri DSM 555

Clostridium lentocellum DSM 5427

Clostridium ljungdahlii DSM 13528

Clostridium novyi NT

Clostridium perfringens ATCC 13124

Clostridium phytofermentans ISDg

Clostridium saccharolyticum WM1

Clostridium sp SY8519

Clostridium sticklandii DSM 519

Clostridium tetani E88

Clostridium thermocellum ATCC 27405

Collimonas fungivorans Ter331

Colwellia psychrerythraea 34H

Comamonas testosteroni CNB 2

Conexibacter woesei DSM 14684

Coprothermobacter proteolyticus DSM 5265

Coraliomargarita akajimensis DSM 45221

Coriobacterium glomerans PW2

Corynebacterium aurimucosum ATCC 700975

Corynebacterium diphtheriae NCTC 13129

Corynebacterium efficiens YS 314

Corynebacterium glutamicum ATCC 13032

Corynebacterium jeikeium K411

Corynebacterium kroppenstedtii DSM 44385

Corynebacterium pseudotuberculosis FRC41

Corynebacterium resistens DSM 45100

Corynebacterium ulcerans 809

Corynebacterium urealyticum DSM 7109

Corynebacterium variabile DSM 44702

Coxiella burnetii RSA 493

Croceibacter atlanticus HTCC2559

Cronobacter sakazakii ATCC BAA 894

Cronobacter turicensis z3032

Cryptobacterium curtum DSM 15641

Cupriavidus metallidurans CH34

Cupriavidus necator N 1

Cupriavidus taiwanensis LMG 19424

cyanobacterium UCYN A

Cyanothece sp ATCC 51142

Cyclobacterium marinum DSM 745

Cytophaga hutchinsonii ATCC 33406

Dechloromonas aromatica RCB

Deferribacter desulfuricans SSM1

Dehalococcoides ethenogenes 195

Dehalococcoides sp BAV1

Dehalogenimonas lykanthroporepellens BL DC 9

Deinococcus deserti VCD115 Deinococcus geothermalis DSM 11300

Deinococcus maricopensis DSM 21211

Deinococcus proteolyticus MRP

Deinococcus radiodurans R1

Delftia acidovorans SPH 1 Delftia sp Cs1 4

Denitrovibrio acetiphilus DSM 12809

Desulfarculus baarsii DSM 2075

Desulfatibacillum alkenivorans AK 01

Desulfitobacterium hafniense Y51

Desulfobacca acetoxidans DSM 11109

Desulfobacterium autotrophicum HRM2

Desulfobulbus propionicus DSM 2032

Desulfococcus oleovorans Hxd3

Desulfohalobium retbaense DSM 5692

Desulfomicrobium baculatum DSM 4028

Desulfotalea psychrophila LSv54

Desulfotomaculum acetoxidans DSM 771

Desulfotomaculum carboxydivorans CO 1 SRB

Desulfotomaculum kuznetsovii DSM 6115

Desulfotomaculum reducens MI 1

Desulfotomaculum ruminis DSM 2154

Desulfovibrio aespoeensis Aspo 2

Desulfovibrio africanus str Walvis Bay

Desulfovibrio alaskensis G20

Desulfovibrio desulfuricans ND132

Desulfovibrio magneticus RS 1

Desulfovibrio salexigens DSM 2638

Desulfovibrio vulgaris RCH1

Desulfurispirillum indicum S5

Desulfurivibrio alkaliphilus AHT2

Desulfurobacterium thermolithotrophum DSM 11699

Desulfurococcus kamchatkensis 1221n

Desulfurococcus mucosus DSM 2162

Dichelobacter nodosus VCS1703A

Dickeya dadantii 3937

Dickeya zeae Ech1591

Dictyoglomus thermophilum H 6 12

Dictyoglomus turgidum DSM 6724

Dinoroseobacter shibae DFL 12

Dyadobacter fermentans DSM 18053

Edwardsiella ictaluri 93 146

Edwardsiella tarda EIB202

Eggerthella lenta DSM 2243

Eggerthella sp YY7918

Ehrlichia canis str Jake

Ehrlichia chaffeensis str Arkansas

Ehrlichia ruminantium str Welgevonden

Elusimicrobium minutum Pei191

Enterobacter aerogenes KCTC 2190

Enterobacter asburiae LF7a

Enterobacter cloacae subsp cloacae ATCC 13047

Enterobacter sp 638 Enterococcus faecalis V583

Erwinia amylovora ATCC 49946

Erwinia billingiae Eb661

Erwinia pyrifoliae DSM 12163

Erwinia sp Ejp617

Erwinia tasmaniensis Et1 99

Erysipelothrix rhusiopathiae

Erythrobacter litoralis HTCC2594

Escherichia coli O157:H7 str Sakai

Escherichia fergusonii ATCC 35469

Ethanoligenens harbinense YUAN 3

Eubacterium eligens ATCC 27750

Eubacterium limosum KIST612

Eubacterium rectale ATCC 33656

Exiguobacterium sibiricum 25515

Exiguobacterium sp AT1b

Ferrimonas balearica DSM 9799

Ferroglobus placidus DSM 10642

Fervidobacterium nodosum Rt17 B1

Fibrobacter succinogenes subsp succinogenes S85

Filifactor alocis ATCC 35896

Finegoldia magna ATCC 29328

Flavobacteriaceae bacterium 3519 10

Flavobacterium branchiophilum FL 15

Flavobacterium columnare ATCC 49512

Flavobacterium johnsoniae UW101

Flavobacterium psychrophilum JIP02 86

Flexistipes sinusarabici DSM 4947

Fluviicola taffensis DSM 16823

Francisella novicida U112

Francisella philomiragia subsp philomiragia ATCC 25017

Francisella sp TX077308

Francisella tularensis subsp holarctica LVS

Frankia alni ACN14a

Frankia sp CcI3

Frankia symbiont of Datisca glomerata

Fusobacterium nucleatum subsp nucleatum ATCC 25586

Gallibacterium anatis UMN179

Gallionella capsiferriformans ES 2 gamma proteobacterium HdN1

Gardnerella vaginalis 409 05

Gemmatimonas aurantiaca T 27

Geobacillus kaustophilus HTA426

Geobacillus sp C56 T3

Geobacillus thermodenitrificans NG80 2

Geobacillus thermoglucosidasius C56 YS93

Geobacillus thermoleovorans CCB US3 UF5

Geobacter bemidjiensis Bem

Geobacter daltonii FRC 32

Geobacter lovleyi SZ

Geobacter metallireducens GS 15

Geobacter sp M18

Geobacter sulfurreducens PCA

Geobacter uraniireducens Rf4

Geodermatophilus obscurus DSM 43160

Glaciecola nitratireducens FR1064

Glaciecola sp 4H 3 7YE 5

Gloeobacter violaceus PCC 7421

Gluconacetobacter diazotrophicus PAl 5

Gluconacetobacter xylinus NBRC 3288

Gluconobacter oxydans 621H

Gordonia bronchialis DSM 43247

Gramella forsetii KT0803

Granulibacter bethesdensis CGDNIH1

Granulicella mallensis MP5ACTX8

Granulicella tundricola

Haemophilus ducreyi 35000HP

Haemophilus influenzae 10810

Haemophilus parainfluenzae T3T1

Haemophilus parasuis SH0165

Haemophilus somnus 129PT

Hahella chejuensis KCTC 2396

Halalkalicoccus jeotgali B3

Halanaerobium hydrogeniformans

Haliangium ochraceum DSM 14365

Haliscomenobacter hydrossis DSM 1100

Haloarcula hispanica ATCC 33960

Haloarcula marismortui ATCC 43049

Halobacterium sp NRC 1

Haloferax volcanii DS2

Halogeometricum borinquense DSM 11551

Halomicrobium mukohataei DSM 12286

Halomonas elongata DSM 2581

halophilic archaeon DL31

Halopiger xanaduensis SH 6

Haloquadratum walsbyi C23

Halorhabdus utahensis DSM 12940

Halorhodospira halophila SL1

Halorubrum lacusprofundi ATCC 49239

Haloterrigena turkmenica DSM 5511

Halothermothrix orenii H 168

Halothiobacillus neapolitanus c2

Helicobacter acinonychis str Sheeba

Helicobacter bizzozeronii CIII 1

Helicobacter felis ATCC 49179

Helicobacter hepaticus ATCC 51449

Helicobacter mustelae 12198

Helicobacter pylori 26695

Heliobacterium modesticaldum Ice1

Herbaspirillum seropedicae SmR1

Herminiimonas arsenicoxydans

Herpetosiphon aurantiacus DSM 785

Hippea maritima DSM 10411

Hirschia baltica ATCC 49814

Hydrogenobacter thermophilus TK 6

Hydrogenobaculum sp Y04AAS1

Hyperthermus butylicus DSM 5456

Hyphomicrobium denitrificans ATCC 51888

Hyphomicrobium sp

Hyphomonas neptunium ATCC 15444

Idiomarina loihiensis L2TR

Ignicoccus hospitalis KIN4 I

Ignisphaera aggregans DSM 17230

Ilyobacter polytropus DSM 2926

Intrasporangium calvum DSM 43043

Isoptericola variabilis 225

Isosphaera pallida ATCC 43644

Jannaschia sp CCS1

Janthinobacterium sp Marseille

Jonesia denitrificans DSM 20603

Kangiella koreensis DSM 16069

Ketogulonicigenium vulgare WSH 001

Kineococcus radiotolerans SRS30216

Kitasatospora setae KM 6054

Klebsiella oxytoca KCTC 1686

Klebsiella pneumoniae 342

Klebsiella variicola At 22

Kocuria rhizophila DC2201

Kosmotoga olearia TBF 1951

Kribbella flavida DSM 17836

Krokinobacter sp 4H 3 7 5

Kyrpidia tusciae DSM 2912

Kytococcus sedentarius DSM 20547

Lacinutrix sp 5H 3 7 4

Lactobacillus acidophilus NCFM

Lactobacillus amylovorus GRL 1112

Lactobacillus brevis ATCC 367

Lactobacillus buchneri NRRL B 30929

Lactobacillus casei ATCC 334

Lactobacillus crispatus ST1

Lactobacillus delbrueckii subsp bulgaricus ATCC 11842

Lactobacillus fermentum IFO 3956

Lactobacillus gasseri ATCC 33323

Lactobacillus helveticus DPC 4571

Lactobacillus johnsonii NCC 533

Lactobacillus kefiranofaciens ZW3

Lactobacillus plantarum subsp plantarum ST III

Lactobacillus reuteri DSM 20016

Lactobacillus rhamnosus ATCC 8530

Lactobacillus ruminis ATCC 27782

Lactobacillus sakei subsp sakei 23K

Lactobacillus salivarius UCC118

Lactobacillus sanfranciscensis TMW 11304

Lactococcus garvieae ATCC 49156

Lactococcus lactis subsp cremoris NZ9000

Laribacter hongkongensis HLHK9

Lawsonia intracellularis PHE MN1 00

Leadbetterella byssophila DSM 17132

Legionella longbeachae NSW150

Legionella pneumophila subsp pneumophila ATCC 43290

Leifsonia xyli subsp xyli str CTCB07

Leptospira biflexa serovar Patoc strain Patoc 1 Ames

Leptospira borgpetersenii serovar Hardjo bovis str L550

Leptospira interrogans serovar Copenhageni str Fiocruz L1 130

Leptothrix cholodnii SP 6

Leptotrichia buccalis C 1013 b

Leuconostoc citreum KM20

Leuconostoc gasicomitatum LMG 18811

Leuconostoc kimchii IMSNU 11154

Leuconostoc mesenteroides subsp mesenteroides ATCC 8293

Leuconostoc sp C2 Listeria innocua Clip11262 Listeria ivanovii

Listeria monocytogenes EGD e

Listeria seeligeri serovar 1 2b str SLCC3954

Listeria welshimeri serovar 6b str SLCC5334

Lysinibacillus sphaericus C3 41

Macrococcus caseolyticus JCSC5402

Magnetococcus marinus MC 1

Magnetospirillum magneticum AMB 1

Mahella australiensis 50 1 BON

Mannheimia succiniciproducens MBEL55E

Maribacter sp HTCC2170

Maricaulis maris MCS10

Marinithermus hydrothermalis DSM 14884

Marinobacter adhaerens HP15

Marinobacter aquaeolei VT8

Marinomonas mediterranea MMB 1

Marinomonas posidonica IVIA Po 181

Marinomonas sp MWYL1

Marivirga tractuosa DSM 4126

Megasphaera elsdenii

Meiothermus ruber DSM 1279

Meiothermus silvanus DSM 9946

Melissococcus plutonius ATCC 35311

Mesoplasma florum L1

Mesorhizobium ciceri biovar biserrulae WSM1271

Mesorhizobium loti MAFF303099

Mesorhizobium opportunistum WSM2075

Metallosphaera cuprina Ar 4

Metallosphaera sedula DSM 5348

Methanobacterium sp AL 21

Methanobrevibacter ruminantium M1

Methanobrevibacter smithii ATCC 35061

Methanocaldococcus fervens AG86

Methanocaldococcus infernus ME

Methanocaldococcus jannaschii DSM 2661

Methanocaldococcus sp FS406 22

Methanocaldococcus vulcanius M7

Methanocella arvoryzae MRE50

Methanocella paludicola SANAE

Methanococcoides burtonii DSM 6242

Methanococcus aeolicus Nankai 3

Methanococcus maripaludis S2

Methanococcus vannielii SB

Methanococcus voltae A3

Methanocorpusculum labreanum Z

Methanoculleus marisnigri JR1

Methanohalobium evestigatum Z 7303

Methanohalophilus mahii DSM 5219

Methanoplanus petrolearius DSM 11571

Methanopyrus kandleri AV19

Methanoregula boonei 6A8

Methanosaeta concilii GP6

Methanosaeta thermophila PT

Methanosalsum zhilinae DSM 4017

Methanosarcina acetivorans C2A

Methanosarcina barkeri str Fusaro

Methanosarcina mazei Go1

Methanosphaera stadtmanae DSM 3091

Methanosphaerula palustris E1 9c

Methanospirillum hungatei JF 1

Methanothermobacter marburgensis str Marburg

Methanothermobacter thermautotrophicus str Delta H

Methanothermococcus okinawensis IH1

Methanothermus fervidus DSM 2088

Methanotorris igneus Kol 5

Methylacidiphilum infernorum V4

Methylibium petroleiphilum PM1

Methylobacillus flagellatus KT

Methylobacterium chloromethanicum CM4

Methylobacterium extorquens AM1

Methylobacterium nodulans ORS 2060

Methylobacterium populi BJ001

Methylobacterium radiotolerans JCM 2831

Methylobacterium sp 4 46

Methylocella silvestris BL2

Methylococcus capsulatus str Bath

Methylomicrobium alcaliphilum

Methylomonas methanica MC09

Methylotenera mobilis JLW8

Methylotenera versatilis 301

Methylovorus glucosetrophus SIP3 4

Methylovorus sp MP688

Micavibrio aeruginosavorus ARL 13

Microbacterium testaceum StLB037

Micrococcus luteus NCTC 2665

Microcystis aeruginosa NIES 843

Microlunatus phosphovorus NM 1

Micromonospora aurantiaca ATCC 27029

Micromonospora sp L5

Mobiluncus curtisii ATCC 43063

Moorella thermoacetica ATCC 39073

Moraxella catarrhalis RH4

Muricauda ruestringensis DSM 13258

Mycobacterium abscessus ATCC 19977

Mycobacterium africanum GM041182

Mycobacterium avium 104

Mycobacterium bovis AF2122 97

Mycobacterium canettii CIPT 140010059

Mycobacterium gilvum PYR GCK

Mycobacterium leprae TN

Mycobacterium marinum M

Mycobacterium rhodesiae NBB3

Mycobacterium smegmatis str MC2 155

Mycobacterium sp JDM601

Mycobacterium tuberculosis H37Rv

Mycobacterium ulcerans Agy99

Mycobacterium vanbaalenii PYR 1

Mycoplasma agalactiae PG2

Mycoplasma arthritidis 158L3 1

Mycoplasma bovis PG45

Mycoplasma capricolum subsp capricolum ATCC 27343

Mycoplasma conjunctivae HRC 581

Mycoplasma crocodyli MP145

Mycoplasma gallisepticum str Rlow

Mycoplasma genitalium G37

Mycoplasma haemocanis str Illinois

Mycoplasma haemofelis str Langford 1

Mycoplasma hominis

Mycoplasma hyopneumoniae 232

Mycoplasma hyorhinis HUB 1

Mycoplasma leachii 99 014 6

Mycoplasma mobile 163K

Mycoplasma mycoides subsp mycoides SC str PG1

Mycoplasma penetrans HF 2

Mycoplasma pneumoniae M129

Mycoplasma pulmonis UAB CTIP

Mycoplasma putrefaciens KS1

Mycoplasma suis str Illinois

Mycoplasma synoviae 53

Myxococcus fulvus HW 1

Myxococcus xanthus DK 1622

Nakamurella multipartita DSM 44233

Nanoarchaeum equitans Kin4 M

Natranaerobius thermophilus JW NM WN LF

Natrialba magadii ATCC 43099

Natronomonas pharaonis DSM 2160

Nautilia profundicola AmH Neisseria gonorrhoeae FA 1090

Neisseria lactamica 020 06

Neisseria meningitidis M01 240149

Neorickettsia risticii str Illinois

Neorickettsia sennetsu str Miyayama

Niastella koreensis GR20 10

Nitratifractor salsuginis DSM 16511

Nitratiruptor sp SB155 2

Nitrobacter hamburgensis X14

Nitrobacter winogradskyi Nb 255

Nitrosococcus halophilus Nc4

Nitrosococcus oceani ATCC 19707

Nitrosococcus watsonii C 113

Nitrosomonas europaea ATCC 19718

Nitrosomonas eutropha C91

Nitrosomonas sp AL212

Nitrosopumilus maritimus SCM1

Nitrosospira multiformis ATCC 25196

Nocardia farcinica IFM 10152

Nocardioides sp JS614

Nocardiopsis dassonvillei subsp dassonvillei DSM 43111

Nostoc azollae 0708

Nostoc punctiforme PCC 73102

Nostoc sp PCC 7120

Novosphingobium aromaticivorans DSM 12444

Novosphingobium sp PP1Y Oceanithermus profundus DSM 14977

Oceanobacillus iheyensis HTE831

Ochrobactrum anthropi ATCC 49188

Odoribacter splanchnicus DSM 20712

Oenococcus oeni PSU 1

Oligotropha carboxidovorans OM5

Olsenella uli DSM 7084

Onion yellows phytoplasma OY M

Opitutus terrae PB90 1

Orientia tsutsugamushi str Boryong

Oscillibacter valericigenes

Owenweeksia hongkongensis DSM 17368

Paenibacillus mucilaginosus KNP414

Paenibacillus polymyxa E681

Paenibacillus sp JDR 2

Paenibacillus terrae HPL 003

Paludibacter propionicigenes WB4

Pantoea ananatis LMG 20103

Pantoea sp At 9b

Pantoea vagans C9 1

Parabacteroides distasonis ATCC 8503

Parachlamydia acanthamoebae UV 7

Paracoccus denitrificans PD1222

Parvibaculum lavamentivorans DS 1

Parvularcula bermudensis HTCC2503

Pasteurella multocida subsp multocida str Pm70

Pectobacterium atrosepticum SCRI1043

Pectobacterium carotovorum subsp carotovorum PC1

Pectobacterium wasabiae WPP163

Pediococcus pentosaceus ATCC 25745

Pedobacter heparinus DSM 2366

Pedobacter saltans DSM 12145

Pelagibacterium halotolerans B2

Pelobacter carbinolicus DSM 2380

Pelobacter propionicus DSM 2379

Pelodictyon phaeoclathratiforme BU 1

Pelotomaculum thermopropionicum SI

Persephonella marina EX H1

Petrotoga mobilis SJ95

Phenylobacterium zucineum HLK1

Photobacterium profundum SS9

Photorhabdus asymbiotica subsp asymbiotica ATCC 43949

Photorhabdus luminescens subsp laumondii TTO1

Picrophilus torridus DSM 9790

Pirellula staleyi DSM 6068

Planctomyces brasiliensis DSM 5305

Planctomyces limnophilus DSM 3776

Polaromonas naphthalenivorans CJ2

Polaromonas sp JS666

Polymorphum gilvum SL003B 26A1

Polynucleobacter necessarius subsp asymbioticus QLW P1DMWA 1

Porphyromonas asaccharolytica DSM 20707

Porphyromonas gingivalis W83

Prevotella denticola F0289

Prevotella melaninogenica ATCC 25845

Prevotella ruminicola 23

Prochlorococcus marinus str MIT 9215

Propionibacterium acnes 6609

Propionibacterium freudenreichii subsp shermanii CIRM BIA1

Prosthecochloris aestuarii DSM 271

Proteus mirabilis HI4320

Pseudoalteromonas atlantica T6c

Pseudoalteromonas haloplanktis TAC125

Pseudoalteromonas sp SM9913

Pseudogulbenkiania sp NH8B

Pseudomonas aeruginosa PAO1

Pseudomonas brassicacearum subsp brassicacearum NFM421

Pseudomonas entomophila L48

Pseudomonas fluorescens F113

Pseudomonas fulva 12 X

Pseudomonas mendocina ymp

Pseudomonas protegens Pf 5

Pseudomonas putida F1

Pseudomonas stutzeri A1501

Pseudomonas syringae pv phaseolicola 1448A

Pseudonocardia dioxanivorans CB1190

Pseudovibrio sp FO BEG1

Pseudoxanthomonas spadix BD a59

Pseudoxanthomonas suwonensis 11 1

Psychrobacter arcticus 273 4

Psychrobacter cryohalolentis K5

Psychrobacter sp PRwf 1

Psychromonas ingrahamii 37

Pusillimonas sp T7 7

Pyrobaculum aerophilum str IM2

Pyrobaculum arsenaticum DSM 13514

Pyrobaculum calidifontis JCM 11548

Pyrobaculum islandicum DSM 4184

Pyrobaculum neutrophilum V24Sta

Pyrobaculum sp 1860

Pyrococcus abyssi GE5

Pyrococcus furiosus DSM 3638

Pyrococcus horikoshii OT3

Pyrococcus sp NA2

Pyrococcus yayanosii CH1

Pyrolobus fumarii 1A

Rahnella sp Y9602

Ralstonia eutropha JMP134

Ralstonia pickettii 12D

Ralstonia solanacearum GMI1000

Ramlibacter tataouinensis TTB310

Renibacterium salmoninarum ATCC 33209

Rhizobium etli CFN 42

Rhizobium leguminosarum bv viciae 3841

Rhodobacter capsulatus SB 1003

Rhodobacter sphaeroides 241

Rhodococcus equi 103S

Rhodococcus erythropolis PR4

Rhodococcus jostii RHA1

Rhodococcus opacus B4

Rhodoferax ferrireducens T118

Rhodomicrobium vannielii ATCC 17100

Rhodopirellula baltica SH 1

Rhodopseudomonas palustris CGA009

Rhodospirillum centenum SW

Rhodospirillum rubrum ATCC 11170

Rhodothermus marinus DSM 4252

Rickettsia africae ESF 5

Rickettsia akari str Hartford

Rickettsia bellii RML369 C

Rickettsia canadensis str McKiel

Rickettsia conorii str Malish 7

Rickettsia felis URRWXCal2

Rickettsia heilongjiangensis 054

Rickettsia japonica YH

Rickettsia massiliae MTU5

Rickettsia peacockii str Rustic

Rickettsia prowazekii str Madrid E

Rickettsia rickettsii str Iowa

Rickettsia sibirica 246

Rickettsia slovaca 13 B

Rickettsia typhi str Wilmington

Riemerella anatipestifer ATCC 11845 DSM 15868

Robiginitalea biformata HTCC2501

Roseburia hominis A2 183

Roseiflexus castenholzii DSM 13941

Roseiflexus sp RS 1

Roseobacter denitrificans OCh 114

Roseobacter litoralis Och 149

Rothia dentocariosa ATCC 17931

Rothia mucilaginosa DY 18

Rubrobacter xylanophilus DSM 9941

Ruegeria pomeroyi DSS 3

Ruegeria sp TM1040

Ruminococcus albus 7

Runella slithyformis DSM 19594

Saccharomonospora viridis DSM 43017

Saccharophagus degradans 2 40

Saccharopolyspora erythraea NRRL 2338

Salinibacter ruber DSM 13855

Salinispora arenicola CNS 205

Salinispora tropica CNB 440

Salmonella bongori NCTC 12419

Salmonella enterica subsp enterica serovar Enteritidis str P125109

Sanguibacter keddieii DSM 10542

Sebaldella termitidis ATCC 33386

Segniliparus rotundus DSM 44985

Selenomonas sputigena ATCC 35185

Serratia plymuthica AS9

Serratia proteamaculans 568 Serratia sp AS12

Serratia symbiotica str Cinara cedri

Shewanella amazonensis SB2B

Shewanella baltica OS155

Shewanella denitrificans OS217

Shewanella frigidimarina NCIMB 400

Shewanella halifaxensis HAW EB4

Shewanella loihica PV 4

Shewanella oneidensis MR 1

Shewanella pealeana ATCC 700345

Shewanella piezotolerans WP3

Shewanella putrefaciens CN 32

Shewanella sediminis HAW EB3

Shewanella sp ANA 3

Shewanella violacea DSS12

Shewanella woodyi ATCC 51908

Shigella boydii Sb227

Shigella dysenteriae Sd197

Shigella flexneri 2a str 301

Shigella sonnei Ss046

Sideroxydans lithotrophicus ES 1

Simkania negevensis Z

Sinorhizobium fredii NGR234

Sinorhizobium medicae WSM419

Sinorhizobium meliloti 1021

Slackia heliotrinireducens DSM 20476

Sodalis glossinidius str morsitans

Sorangium cellulosum So ce56

Sphaerobacter thermophilus DSM 20745

Sphaerochaeta coccoides DSM 17374

Sphaerochaeta globus str Buddy

Sphaerochaeta pleomorpha str Grapes

Sphingobacterium sp 21

Sphingobium chlorophenolicum L 1

Sphingobium japonicum UT26S

Sphingobium sp SYK 6

Sphingomonas wittichii RW1

Sphingopyxis alaskensis RB2256

Spirochaeta caldaria DSM 7334

Spirochaeta smaragdinae DSM 11293

Spirochaeta thermophila DSM 6192

Spirosoma linguale DSM 74

Stackebrandtia nassauensis DSM 44728

Staphylococcus aureus subsp aureus JH1

Staphylococcus carnosus subsp carnosus TM300

Staphylococcus epidermidis ATCC 12228

Staphylococcus haemolyticus JCSC1435

Staphylococcus lugdunensis HKU09 01

Staphylococcus pseudintermedius HKU10 03

Staphylothermus hellenicus DSM 12710

Staphylothermus marinus F1

Starkeya novella DSM 506

Stenotrophomonas maltophilia K279a

Stigmatella aurantiaca DW4 3 1

Streptobacillus moniliformis DSM 12112

Streptococcus agalactiae 2603V R

Streptococcus dysgalactiae subsp equisimilis ATCC 12394

Streptococcus equi subspecies zooepidemicus

Streptococcus gallolyticus subsp gallolyticus ATCC BAA 2069

Streptococcus gordonii str Challis substr CH1

Streptococcus macedonicus ACA DC 198

Streptococcus mitis B6

Streptococcus mutans NN2025

Streptococcus oralis Uo5

Streptococcus parasanguinis ATCC 15912

Streptococcus parauberis KCTC 11537

Streptococcus pasteurianus ATCC 43144

Streptococcus pneumoniae ST556

Streptococcus pseudopneumoniae IS7493

Streptococcus pyogenes M1 GAS

Streptococcus salivarius JIM8780

Streptococcus sanguinis SK36

Streptococcus suis 05ZYH33

Streptococcus thermophilus CNRZ1066

Streptococcus uberis 0140J

Streptomyces avermitilis MA 4680

Streptomyces bingchenggensis BCW 1

Streptomyces cattleya NRRL 8057 DSM 46488

Streptomyces coelicolor A32

Streptomyces flavogriseus ATCC 33331

Streptomyces griseus subsp griseus NBRC 13350

Streptomyces scabiei 8722

Streptomyces sp SirexAA E

Streptomyces violaceusniger Tu 4113

Streptosporangium roseum DSM 43021

Sulfobacillus acidophilus DSM 10332

Sulfolobus acidocaldarius DSM 639

Sulfolobus islandicus M1425

Sulfolobus solfataricus P2

Sulfolobus tokodaii str 7

Sulfuricurvum kujiense DSM 16994

Sulfurihydrogenibium azorense Az Fu1

Sulfurihydrogenibium sp YO3AOP1

Sulfurimonas autotrophica DSM 16294

Sulfurimonas denitrificans DSM 1251

Sulfurospirillum deleyianum DSM 6946

Sulfurovum sp NBC37 1

Symbiobacterium thermophilum IAM 14863

Synechococcus elongatus PCC 6301

Synechococcus sp CC9605

Synechocystis sp PCC 6803

Syntrophobacter fumaroxidans MPOB

Syntrophobotulus glycolicus DSM 8271

Syntrophomonas wolfei subsp wolfei str Goettingen

Syntrophothermus lipocalidus DSM 12680

Syntrophus aciditrophicus SB

Tannerella forsythia ATCC 43037

Taylorella equigenitalis MCE9

Tepidanaerobacter acetatoxydans Re1

Teredinibacter turnerae T7901

Terriglobus saanensis SP1PR4

Tetragenococcus halophilus

Thauera sp MZ1T

Thermaerobacter marianensis DSM 12885

Thermanaerovibrio acidaminovorans DSM 6589

Thermincola potens JR

Thermoanaerobacter brockii subsp finnii Ako 1

Thermoanaerobacter italicus Ab9

Thermoanaerobacterium thermosaccharolyticum DSM 571

Thermoanaerobacterium xylanolyticum LX 11

Thermoanaerobacter mathranii subsp mathranii str A3

Thermoanaerobacter pseudethanolicus ATCC 33223

Thermoanaerobacter sp X513

Thermoanaerobacter tengcongensis MB4

Thermoanaerobacter wiegelii Rt8B1

Thermobaculum terrenum ATCC BAA 798

Thermobifida fusca YX

Thermobispora bispora DSM 43833

Thermococcus barophilus MP

Thermococcus gammatolerans EJ3

Thermococcus kodakarensis KOD1

Thermococcus onnurineus NA1

Thermococcus sibiricus MM 739

Thermococcus sp 4557

Thermocrinis albus DSM 14484

Thermodesulfatator indicus DSM 15286

Thermodesulfobacterium sp OPB45

Thermodesulfobium narugense DSM 14796

Thermodesulfovibrio yellowstonii DSM 11347

Thermofilum pendens Hrk 5

Thermomicrobium roseum DSM 5159

Thermomonospora curvata DSM 43183

Thermoplasma acidophilum DSM 1728

Thermoplasma volcanium GSS1

Thermoproteus uzoniensis 76820

Thermosediminibacter oceani DSM 16646

Thermosipho africanus TCF52B

Thermosipho melanesiensis BI429

Thermosphaera aggregans DSM 11486

Thermosynechococcus elongatus BP 1

Thermotoga lettingae TMO

Thermotoga maritima MSB8

Thermotoga naphthophila RKU 10

Thermotoga neapolitana DSM 4359

Thermotoga petrophila RKU 1

Thermotoga sp RQ2

Thermotoga thermarum DSM 5069

Thermovibrio ammonificans HB 1

Thermovirga lienii DSM 17291

Thermus scotoductus SA 01

Thermus sp CCB US3 UF1

Thermus thermophilus HB27

Thioalkalimicrobium cyclicum ALM1

Thioalkalivibrio sp K90mix

Thioalkalivibrio sulfidophilus HL EbGr7

Thiobacillus denitrificans ATCC 25259

Thiomicrospira crunogena XCL 2

Thiomonas intermedia K12

Tolumonas auensis DSM 9187

Treponema azotonutricium ZAS 9

Treponema brennaborense DSM 12168

Treponema denticola ATCC 35405

Treponema pallidum subsp pallidum str Nichols

Treponema paraluiscuniculi Cuniculi A

Treponema primitia ZAS 2

Treponema succinifaciens DSM 2489

Trichodesmium erythraeum IMS101

Tropheryma whipplei str Twist

Truepera radiovictrix DSM 17093

Tsukamurella paurometabola DSM 20162

Ureaplasma parvum serovar 3 str ATCC 27815

Ureaplasma urealyticum serovar 10 str ATCC 33699

Variovorax paradoxus S110

Veillonella parvula DSM 2008

Verminephrobacter eiseniae EF01 2

Verrucosispora maris AB 18 032

Vibrio anguillarum 775

Vibrio cholerae O1 biovar El Tor str N16961

Vibrio fischeri ES114

Vibrio furnissii NCTC 11218

Vibrio harveyi ATCC BAA 1116

Vibrio parahaemolyticus RIMD 2210633

Vibrio sp EJY3

Vibrio splendidus LGP32

Vibrio vulnificus CMCP6

Vulcanisaeta distributa DSM 14429

Vulcanisaeta moutnovskia 768 28

Waddlia chondrophila WSU 86 1044

Weeksella virosa DSM 16922

Weissella koreensis KACC 15510

Wigglesworthia glossinidia endosymbiont of Glossina brevipalpis

Wolbachia endosymbiont of Culex quinquefasciatus Pel Wolbachia sp wRi

Wolinella succinogenes DSM 1740

Xanthobacter autotrophicus Py2

Xanthomonas albilineans GPE PC73

Xanthomonas axonopodis pv citri str 306

Xanthomonas campestris pv campestris str 8004

Xanthomonas oryzae pv oryzae KACC 10331

Xenorhabdus bovienii SS 2004

Xenorhabdus nematophila ATCC 19061

Xylanimonas cellulosilytica DSM 15894

Xylella fastidiosa 9a5c

Yersinia enterocolitica subsp enterocolitica 8081

Yersinia pestis CO92

Yersinia pseudotuberculosis IP 32953

Zobellia galactanivorans

Zunongwangia profunda SM A87

Zymomonas mobilis subsp mobilis ATCC 10988

